# Lupus Immune Complexes Drive Distinct Pro-Inflammatory Monocyte and Macrophage Populations Independent of Type I Interferon

**DOI:** 10.64898/2026.01.21.700902

**Authors:** Lais Osmani, Min Shin, Sang Jin Lee, Helen Cai, Won Jae Seong, Hyoungsu Kim, Jongjin Yoo, Minhyung Kim, William Bracamonte, Mario Felix, Jong Gyun Ahn, Hong-Jai Park, Junghee J Shin, Serhan Unlu, Jennefer Par-Young, Edward Doherty, Jiaye Chen, Mei X Dong, Fotios Koumpouras, Jose L Gomez, Naftali Kaminski, Richard Bucala, Sungyong You, Insoo Kang

## Abstract

Systemic lupus erythematosus (SLE or lupus) is an autoimmune disease characterized by anti-nuclear antibody (ANA) production and inflammation, though the mechanisms by which ANAs induce inflammation and tissue injury are incompletely understood. Here, we identified distinct subsets of mononuclear phagocytes (MPs), including monocytes (Mo) and macrophages (MΦ), driven by lupus immune complex (IC), comprised of ANAs and their target antigens. scRNA-seq of human Mo incubated with U1-snRNP (snRNP) lupus IC revealed expansion of distinct pro-inflammatory Mo subsets with upregulation of inflammatory genes including those encoding cytokines, NLRP3, and transcription factors. These transcriptomic changes strongly correlated with protein expression, as determined by proteomic analysis. Mo developed similar pro-inflammatory transcriptomic changes in response to other lupus ICs containing anti-dsDNA and Ro60 antibodies. Interrogation of scRNA-seq datasets from the skin, kidney, and peripheral blood of lupus patients revealed the presence and expansion of pro-inflammatory Mo and MΦ populations exhibiting transcriptomic signatures similar to those observed in lupus IC-stimulated Mo. Some of these cells expressing the snRNP IC gene signature exhibited low expression of the type I IFN signature, suggesting that lupus IC and type I IFN signaling may independently affect Mo subsets. In lupus nephritis, infiltration of CD68+ MΦ expressing NLRP3 was associated with treatment outcomes. Inhibiting activation of the transcription factor ETS2, a master regulator of Mo/MΦ-driven inflammation, attenuated lupus IC-induced activation of pro-inflammatory Mo. Collectively, these findings provide novel insights into the role of lupus IC-driven pro-inflammatory MPs in the pathogenesis of SLE and highlight their relevance as therapeutic targets.

**Significance Statement:** Systemic lupus erythematosus (SLE or lupus) is a multi-systemic autoimmune inflammatory disease characterized by anti-nuclear antibody (ANA) production. Lupus immune complex (IC), consisting of ANAs and their target antigens, likely play a critical role in the pathogenesis of lupus through activation of mononuclear phagocytes (MPs), including monocytes (Mo) and macrophages (MΦ), which can produce an array of inflammatory molecules. Using transcriptomic and proteomic analyses, our study identified distinct pro-inflammatory Mo populations driven by lupus IC, along with expansion of similar pro-inflammatory Mo and MΦ subsets in the skin, kidneys, and peripheral blood of lupus patients. These findings provide novel insights into the pathogenic role of lupus IC-driven pro-inflammatory MPs and support a scientific rationale for therapeutically targeting these cells.

**Graphical Abstract:** 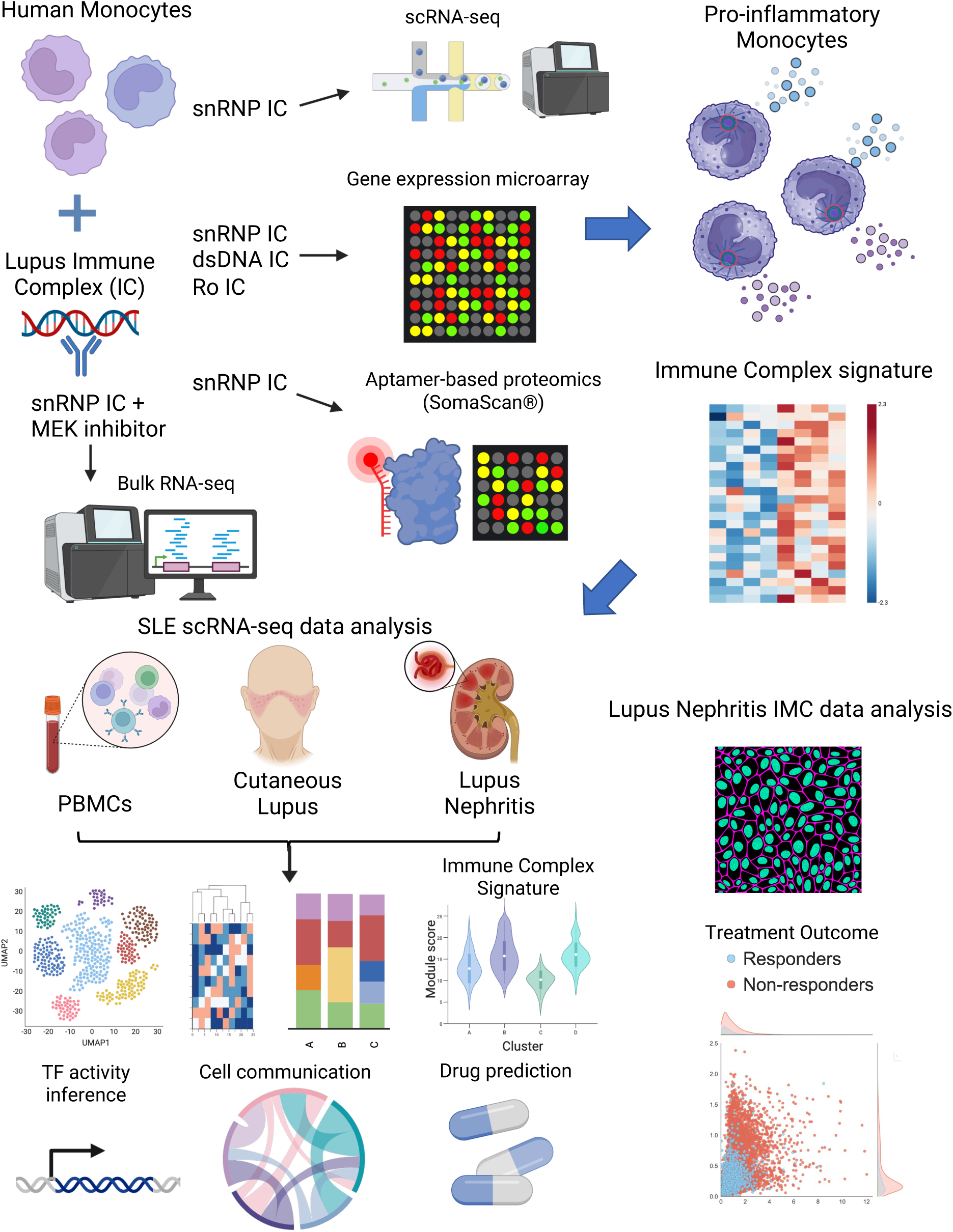

## Introduction

Systemic lupus erythematosus (SLE or lupus) is a multi-systemic autoimmune-mediated inflammatory disease of unknown etiology involving multiple organs. The pathologic features of SLE are thought to involve alterations in the immune response to autoantigens, especially ones in the nucleus, leading to autoantibody production with subsequent inflammation and tissue injury (*1*). In addition to adaptive immunity including B and T cells, the roles of the innate immune system in SLE have been emerging recently. These include neutrophil extracellular trap (NET) formation, clearance of apoptotic debris, production of cytotoxic molecules, cytokines (*e.g.,* type I IFN) and chemokines, and activation of adaptive immune cells (*1*). Lupus patients exhibit diverse clinical manifestations, which range from mild joint and skin disease to serious deep organ involvement such as nephritis, suggesting the involvement of diverse pathogenic mechanisms. Dissecting the latter point may lead to therapies that target the dominant clinical manifestations in individual lupus patients rather than utilizing global immunosuppressive medications such as glucocorticoids and cytotoxic agents with significant adverse effects.

Mononuclear phagocytes (MPs), including monocytes (Mo) and macrophages (MΦ), are main components of the innate immune system (*2*). Recent studies including those from single cell RNA sequencing (scRNA-seq) analysis have revealed more complex heterogeneity in Mo and MΦ than originally appreciated (*3, 4*). The functional characteristics of Mo and MΦ, including phagocytosis, antigen presentation, cytokine production, and tissue infiltration, support the involvement of Mo and MΦ in the pathogenesis of autoimmune disease (*5*). Lupus immune complexes (IC), including U1-snRNP/anti-U1-snRNP antibody (Ab) and dsDNA/anti-dsDNA Ab (referred to as snRNP IC and dsDNA IC, respectively), can activate human Mo via triggering endosomal Toll-like receptors (TLRs) 7, 8, and 9 with self-nucleic acids and the NLRP3 inflammasome, leading to the production of the pro-inflammatory cytokines IL-1β and IL-18 (*6, 7*). Such NLRP3 inflammasome activation is enhanced by the cytokine macrophage migration inhibitory factor (MIF), which is also a susceptibility gene for severe lupus, released from Mo upon lupus IC stimulation (*8*). A previous study reported the possible roles of TLR7 and TLR9 in SLE using genetically modified lupus-prone mice (*9*). Another study demonstrated the potential roles of Mo and TLRs in the development of lupus nephritis in murine lupus models by identifying TLR-activated patrolling Mo as a contributor to glomerular inflammation and kidney injury (*10*). Moreover, a TLR7 gain-of-function genetic variation was shown to cause human lupus, supporting the importance of TLR7 and its guanosine-containing self-ligands, such as U1-snRNP, in the pathogenesis of SLE (*11*).

This study aimed to explore the functional heterogeneity of Mo and MΦ in the pathogenesis of SLE, focusing on activation driven by lupus IC. We characterized transcriptomic and proteomic changes in distinct lupus IC activated Mo and identified corresponding pro-inflammatory Mo and MΦ populations in the skin, kidneys, and peripheral blood of lupus patients through analysis of scRNA-seq, proteomic, and Imaging Mass Cytometry (IMC) data. Our study identified distinct pro-inflammatory Mo populations driven by lupus IC-mediated activation along with expansion of similar pro-inflammatory Mo and MΦ subsets in the skin, kidneys, and peripheral blood of lupus patients that likely serve as key mediators in inflammation in SLE.

## Results

### U1-snRNP immune complex (snRNP IC) induces expansion of pro-inflammatory subsets of human Mo as determined by scRNA-seq analysis

We previously reported that human Mo activated with lupus IC, such as snRNP IC, produced high levels of multiple cytokines (*e.g.,* IL-1β, TNF-⍺, IL-6, IL-18), whereas anti-snRNP Ab+ serum or snRNP alone hardly induced cytokine production by Mo (*6, 7*). To explore the global transcriptomic changes in such activated Mo at the single cell level, untouched Mo (CD14^+^CD16^-^) from healthy individuals were incubated with or without snRNP IC followed by scRNA-seq analysis. We identified eight distinct clusters from 3,040 Mo, which were differentially distributed in Mo stimulated with or without snRNP IC (Fig. 1A-B). Based on differentially expressed genes (DEGs), these clusters were assigned to the following eight distinct Mo subsets (Fig. 1A and Fig. S1A): S100 pro-inflammatory Mo, S100^low^ pro-inflammatory Mo, CYP1B1/C3AR1 Mo, S100/NLRP12/CCR2 Mo, GZMB/NLRP7 Mo, PADI2/HLA Mo, CLEC10A/CCR7 Mo, and CD34/SPINK2 Mo. Most notably, the S100 pro-inflammatory Mo, which expressed genes encoding pro-inflammatory cytokines and chemokines (i.e. S100A8/9, TNF, IL-1β, IL-18, IL-6, CCL3, CCL4, and CCL20) and NLRP3, expanded from 2.5% to 62.2% upon snRNP IC stimulation (Fig. 1C). Among unstimulated Mo, the S100/NLRP12/CCR2 Mo defined by high expression of *S100A8/9* encoding calgranulin A and B, heterodimers of which form calprotectin, was the most abundant Mo subset, and substantially contracted with snRNP IC stimulation. While both S100 pro-inflammatory Mo and S100/NLRP12/CCR2 Mo expressed high levels of S*100A8* and *S100A9*, other inflammatory genes were minimally expressed by S100/NLRP12/CCR2 Mo (Fig. 1A heatmap and Fig. S1A). NLRP12 was shown to be a negative regulator of Mo activation and cytokine production via inhibition of NF-kB activation (*12*). Thus, lupus IC may activate S100/NLRP12/CCR2 Mo resulting in upregulation of pro-inflammatory genes and transformation into a pro-inflammatory phenotype. Indeed, the S100 pro-inflammatory Mo exhibited the highest expression of the cytokine-cytokine receptor interaction and the Toll-like receptor (TLR) signaling pathway gene sets (Fig. 1D and Fig. S1B). There was a modest expansion of the S100^low^ pro-inflammatory Mo (Fig. 1C) which also expressed pro-inflammatory genes (Fig. 1A heatmap and Fig. S1A). The CLEC10A/CCR7, GZMB/NLRP7 and PADI2/HLA Mo subsets exhibited significantly higher expression of antigen processing and presentation gene sets compared to other subsets (Fig. 1D).

**Figure 1.**
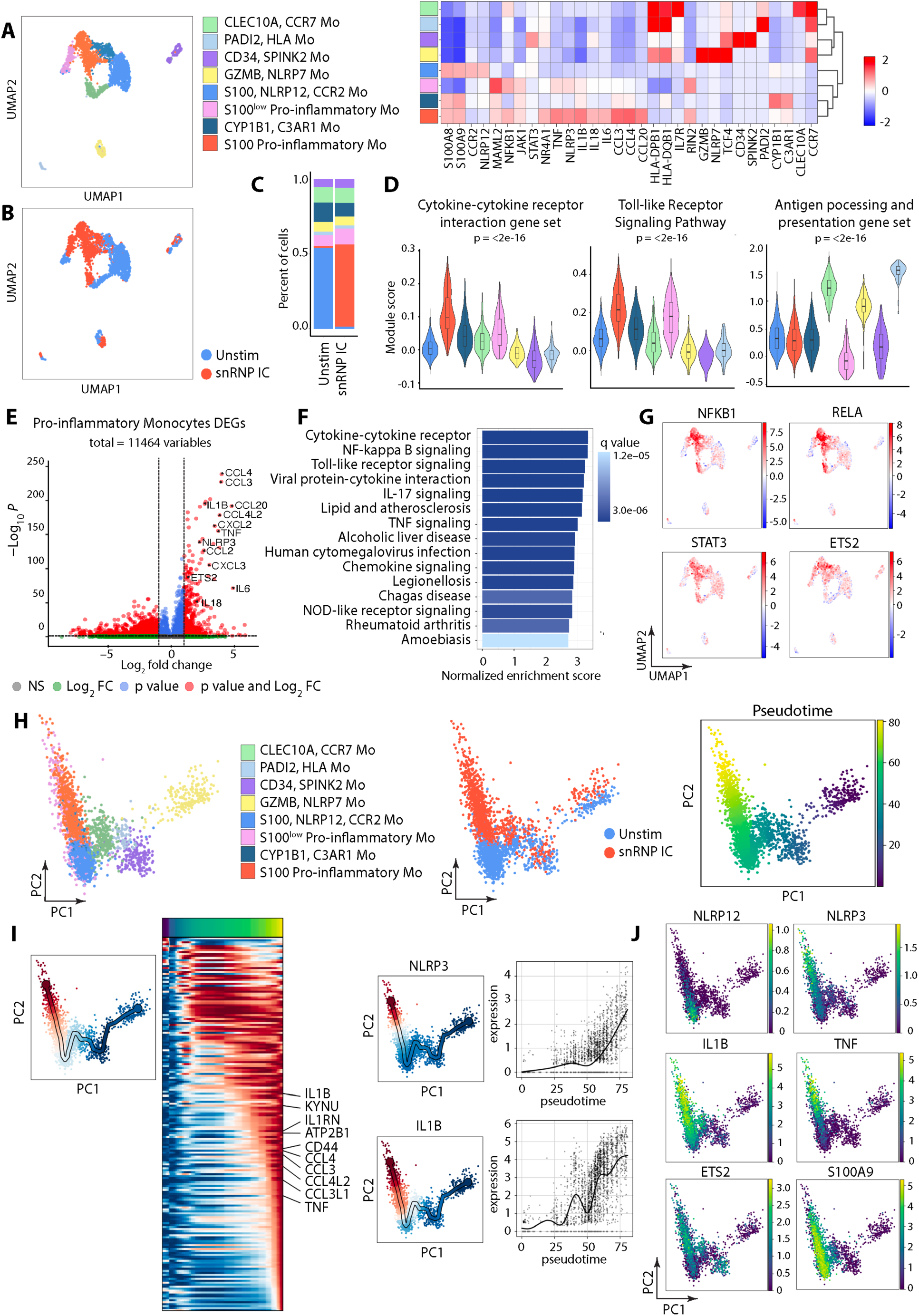
Single-cell transcriptomic profiling of snRNP immune complex (IC) stimulated monocytes (Mo) identifies pro-inflammatory Mo subsets. Untouched Mo (CD14^+^CD16^-^) isolated from the PBMCs of healthy adult subjects (n = 2) were incubated for 2 hours with or without U1-snRNP and pooled anti-U1-snRNP antibody (Ab) positive serum (snRNP IC) followed by scRNA-seq analysis. (A) UMAP visualization of Mo clusters and scaled heatmap showing mean expression of selected differentially expressed genes (DEGs) used for cluster annotation. (B) UMAP colored by condition (blue = unstimulated Mo, red = snRNP IC stimulated Mo). (C) Stacked bar plot displaying the relative proportions of Mo clusters across the two conditions. (D) Violin plots of module scores for expression of indicated gene sets. Statistical comparisons were performed using the Kruskal-Wallis test. (E) Volcano plot of DEGs from S100 pro-inflammatory Mo versus other Mo clusters (adjusted *P* < 0.05, |log_2_fold change| > 0.5). (F) Bar plot showing the top 15 enriched KEGG pathways by Gene Set Enrichment Analysis (GSEA), ranked by normalized enrichment score. (G) Feature plots illustrating inferred transcription factor activity of indicated transcription factors. (H) PCA plot of Mo clusters, PCA plot colored by condition (blue = unstimulated Mo, red = snRNP IC stimulated Mo), and PCA plot colored by pseudotime trajectory. (I) Heatmap of gene expression dynamics along the terminal end of pseudotime, as indicated by accompanying PCA plot and expression of *NLRP3* and *IL1B* along pseudotime progression. (J) Feature plots of *NLRP12*, *NLRP3*, *IL1B*, *TNF*, ETS2 and *S100A9 expression*.

We further characterized S100 pro-inflammatory Mo given the significant expansion of this subset following snRNP IC stimulation. Differential gene expression analysis between S100 pro-inflammatory Mo and other Mo subsets demonstrated significant upregulation of pro-inflammatory genes (Fig. 1E), including the top marker genes shown in Fig 1A (i.e., *IL1B, IL18, TNF, CCL3, CCL4 and NLRP3*), suggesting enhanced inflammatory function of S100 pro-inflammatory Mo. Gene Set Enrichment Analysis (GSEA) of KEGG pathways revealed significant enrichment of NOD-like receptor signaling pathway, TLR signaling, NF-kappa B signaling pathway, TNF signaling pathway, chemokine signaling pathway and cytokine-cytokine receptor interaction (Fig. 1F). We performed transcription factor (TF) activity inference using decoupleR (*13*) and found highest activity of NFKB1, RELA (p65), STAT3, and ETS2 within the S100 pro-inflammatory Mo (Fig. 1G). Notably, ETS2 was recently identified as a master regulator of inflammatory MΦ (*14*). These findings are in line with our published studies implicating NF-kB and TLRs in the activation of Mo in response to lupus IC, including snRNP IC (6–8).

### Pseudotime trajectory analysis reveals transition to pro-inflammatory Mo from S100/NLRP12/CCR2 Mo in response to snRNP IC

We performed trajectory analysis using scFates (*15*) to infer potential transitions between subsets in response to snRNP IC stimulation. Trajectory analysis identified S100 pro-inflammatory and S100^low^ pro-inflammatory Mo as the most terminal subsets along the pseudotime trajectory (Fig. 1H). These Mo subsets appeared to arise from S100/NLRP12/CCR2 Mo upon snRNP IC stimulation. With pseudotime progression, S100 pro-inflammatory and S100^low^ pro-inflammatory Mo demonstrated increased expression of multiple pro-inflammatory genes (Fig. 1I and Fig. S2). The transition from S100/NLRP12/CCR2 Mo to pro-inflammatory Mo was accompanied by decreased expression of *NLRP12* and increased expression of *NLRP3* and other inflammatory genes (Fig. 1J). This finding is in line with the reciprocal changes in the proportions of S100 pro-inflammatory and S100/NLRP12/CCR2 Mo with snRNP IC stimulation (Fig 1C).

### Mo exhibit highly overlapping transcriptomic changes in response to snRNP, Ro60, and dsDNA IC

We assessed whether a similar pro-inflammatory gene signature identified by our scRNA-seq analysis could be detected in Mo stimulated with different types of lupus-associated IC, including dsDNA IC, Ro60/anti-Ro60 (Ro) IC, and snRNP IC. Our microarray analysis of Mo incubated with the three types of IC identified sets of DEGs compared to Mo incubated without IC (Fig. 2A). Substantial overlap of DEGs was observed in Mo stimulated with the three types of IC (Fig 2A). Functional enrichment analysis of upregulated DEGs revealed enrichment of immune-related pathways including NOD-like receptor signaling pathway, NF-kappa B signaling pathway, TNF signaling pathway, and TLR signaling pathway in Mo stimulated with all three types of IC (Fig. 2B). These pathways overlap with the enriched pathways found in S100 pro-inflammatory Mo by scRNA-seq analysis of Mo stimulated with snRNP IC (Fig. 1F). Correlation analysis of DEGs in Mo stimulated with the three types of lupus IC demonstrated significant correlations (Fig. 2C). We explored whether the gene signatures derived from upregulated DEGs in bulk Mo stimulated with the three distinct types of lupus IC could be identified within the Mo subsets stimulated with snRNP IC in our scRNA-seq data. Expression of upregulated DEGs from bulk Mo stimulated with snRNP IC, Ro IC and dsDNA IC was highest in S100 pro-inflammatory Mo followed by S100^low^ pro-inflammatory Mo (Fig. 2D). Overall, these findings demonstrate that lupus-associated dsDNA, Ro, and snRNP IC induce largely similar responses in inflammatory subsets of Mo.

**Figure 2.**
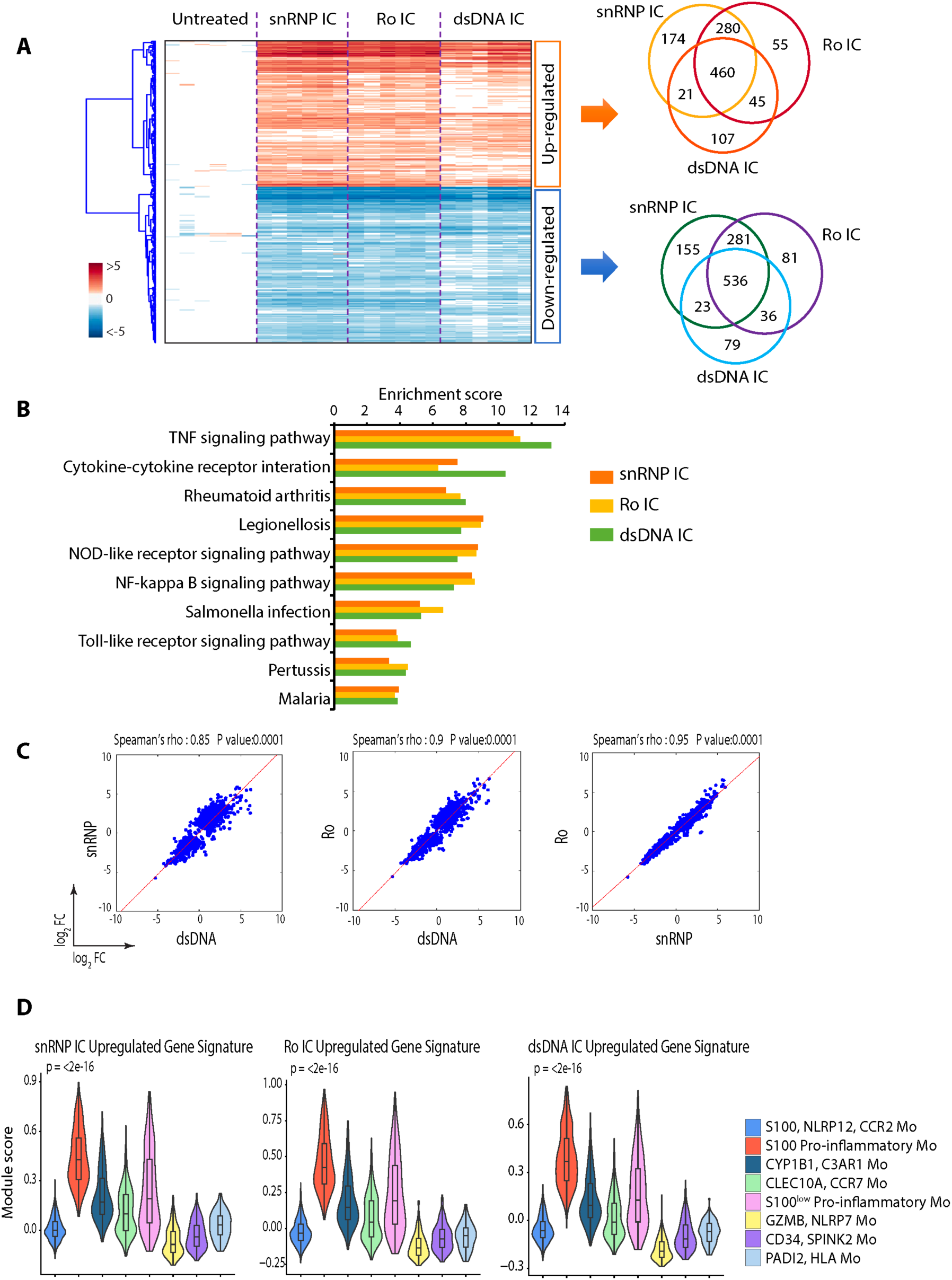
Monocytes (Mo) exhibit highly overlapping transcriptomic changes in response to snRNP, Ro60, and dsDNA immune complexes (IC). Microarray analysis of untouched Mo (CD14^+^CD16^-^) isolated from the PBMCs of healthy adult subjects (n = 6) were incubated for 3 hours with or without U1-snRNP and pooled anti-U1-snRNP antibody (Ab) positive serum (snRNP IC), human genomic dsDNA (5 μg/ml) and pooled anti-dsDNA Ab+ serum (dsDNA IC), or Ro60 (5 μg/ml) and pooled anti-Ro60 Ab+ serum (Ro IC). (A) Heatmap and Venn diagrams of upregulated and downregulated genes in response to indicated IC stimulation. (B) Top KEGG pathways enriched from upregulated genes. (C) Pairwise Spearman correlation scatter plots comparing differentially expressed genes (DEGs) by Mo simulated with indicated IC. (D) Violin plots of module scores in scRNA-seq data of Figure 1 showing expression of upregulated genes identified from microarray analysis of Mo stimulated with indicated IC. Statistical comparisons of module scores were performed using the Kruskal-Wallis test.

### IC signature genes and their corresponding proteins highly correlate in Mo stimulated with snRNP IC

We investigated whether our gene expression findings translated into protein production by performing high-plex SomaScan® proteomics analysis of the culture supernatant of Mo stimulated with or without snRNP IC. This analysis identified many differentially expressed proteins (DEPs), including cytokines, chemokines, and other inflammation associated proteins (Fig. 3A). Enriched KEGG pathways among upregulated proteins include cytokine-cytokine receptor interaction, chemokine signaling, TNF signaling, NOD-like receptor signaling, TLR signaling, and NF-kappa B signaling pathways (Fig. 3B). This finding is in line with the results of our transcriptomic analyses (Fig. 1F and Fig. 2B). Our multiplex immunoassay on the same culture supernatant of Mo corroborated our proteomic findings and also identified increased production of B-cell activating factor (BAFF or TNFSF13B) from Mo stimulated with snRNP IC. (Fig. 3C). BAFF is crucial for B cell survival, maturation, Ab production and class switching (*16*), and anti-BAFF Ab therapy has been approved by the FDA for treating lupus.

**Figure 3.**
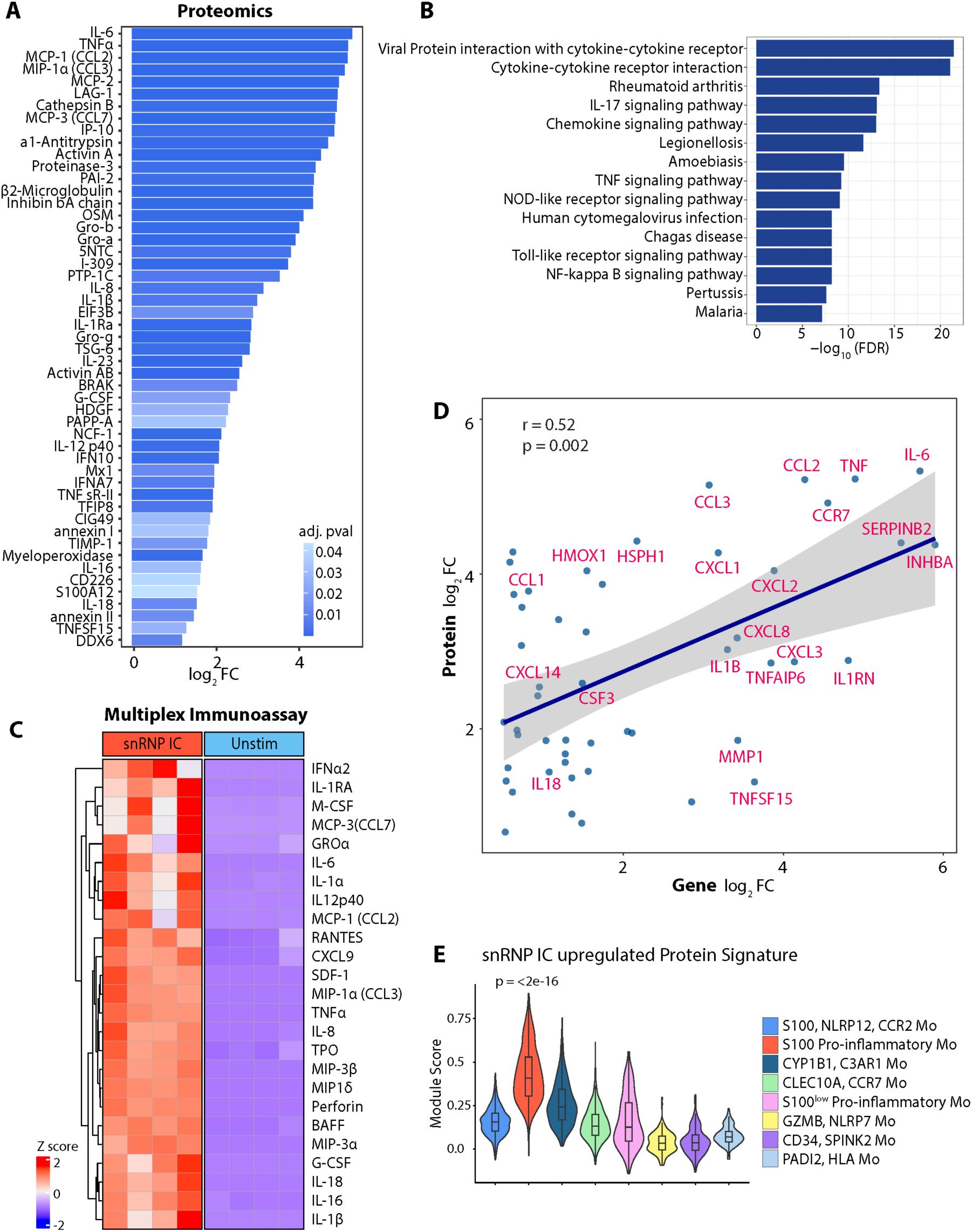
Transcriptomic and proteomic changes highly correlate in monocytes (Mo) stimulated with snRNP immune complex (IC). Culture supernatants of untouched Mo (CD14^+^CD16^-^) which were incubated for 14 hours with or without U1-snRNP and pooled anti-U1-snRNP antibody (Ab) positive serum (snRNP IC) were analyzed using SomaScan® (n = 5) (A, B, D, and E) and cytokine/chemokine multiplex (n = 4) (C) assays. (A) Bar plot showing upregulated proteins in snRNP IC-stimulated Mo compared to unstimulated Mo (adjusted *P* < 0.05). (B) Top 15 KEGG pathways enriched from upregulated differentially expressed proteins (DEPs). (C) Scaled heatmap of concentrations of selected proteins measured by cytokine/chemokine multiplex assay. (D) Spearman correlation scatter plot comparing upregulated genes (log_2_fold change > 0.5) from microarray analysis of Figure 2 and upregulated proteins (log_2_fold change > 0.5) in response to snRNP IC. (E) Violin plot of module scores in scRNA-seq data of Figure 1 for expression of genes corresponding to upregulated DEPs in response to snRNP IC stimulation. Statistical comparisons of module scores were performed using the Kruskal-Wallis test.

Spearman correlation analysis revealed a significant positive correlation between upregulated genes from microarray analysis and upregulated proteins in response to snRNP IC stimulation (Fig. 3D). Many of these proteins are cytokines, chemokines or inflammation-associated. Using the genes corresponding to the upregulated proteins, we created a snRNP IC-upregulated protein signature gene set. Expression of this upregulated protein signature was highest in S100 pro-inflammatory Mo compared to other Mo subsets (Fig. 3E).

### Pro-inflammatory MPs with lupus IC signature are present in active cutaneous lupus lesions as determined by scRNA-seq analysis

To assess the potential pathologic implications of lupus IC *in vivo*, we interrogated whether Mo and/or MΦ, which can be derived from activated Mo, in active cutaneous lupus exhibit transcriptomic features similar to pro-inflammatory Mo subsets activated *ex vivo* with lupus IC using publicly available scRNA-seq data (GSE186476) (*17*). This scRNA-seq dataset consisting of biopsies from 7 active cutaneous lupus lesions and 7 normal skin tissue specimens was normalized, integrated, and analyzed. A subset of MPs, including MΦ and dendritic cells (DCs), was identified based on expression of *LYZ, CD14,* and *FCGR3A* (n = 591 cells). Clustering of these cells yielded six clusters, which were assigned based on their DEGs and known canonical markers (Fig. 4A-B): S100 pro-inflammatory MΦ, interferon-stimulated genes (ISG) pro-inflammatory MPs, Mo derived dendritic cells (moDCs), Langerhans cells, S100 MΦ, and C1Q/FOLR2/APOE MΦ. Active cutaneous lupus skin exhibited a notable expansion of S100 pro-inflammatory MΦ and ISG pro-inflammatory MPs as well as contraction of S100 MΦ compared to healthy control skin (Fig. 4C). The S100 pro-inflammatory MΦ and ISG pro-inflammatory MPs demonstrated highest expression of pro-inflammatory cytokines and chemokines including *TNF*, *IL1A, IL1B, IL6, CCL2, CCL3*, and *CCL4* (Fig. 4A heatmap). Both S100 pro-inflammatory MΦ and ISG pro-inflammatory MPs demonstrated the highest expression of the snRNP IC gene and protein signatures identified by our scRNA-seq and proteomics analysis of snRNP IC-stimulated Mo (Fig. 4D). However, only ISG pro-inflammatory MPs exhibited high expression of type I IFN inducible genes (Fig. 4D). Similar findings were observed for the expression of dsDNA and Ro IC signatures from our microarray analysis (Fig. S3A). These findings suggest that the differentiation of S100 pro-inflammatory MΦ is dependent on activation by lupus IC, while the differentiation of ISG proinflammatory MPs is driven by both lupus IC and type I IFN. ISG pro-inflammatory MPs and S100 pro-inflammatory MΦ also demonstrated the highest expression of KEGG TLR, NOD-like receptor signaling, and cytokine-cytokine receptor interaction pathways (Fig. S3B).

**Figure 4.**
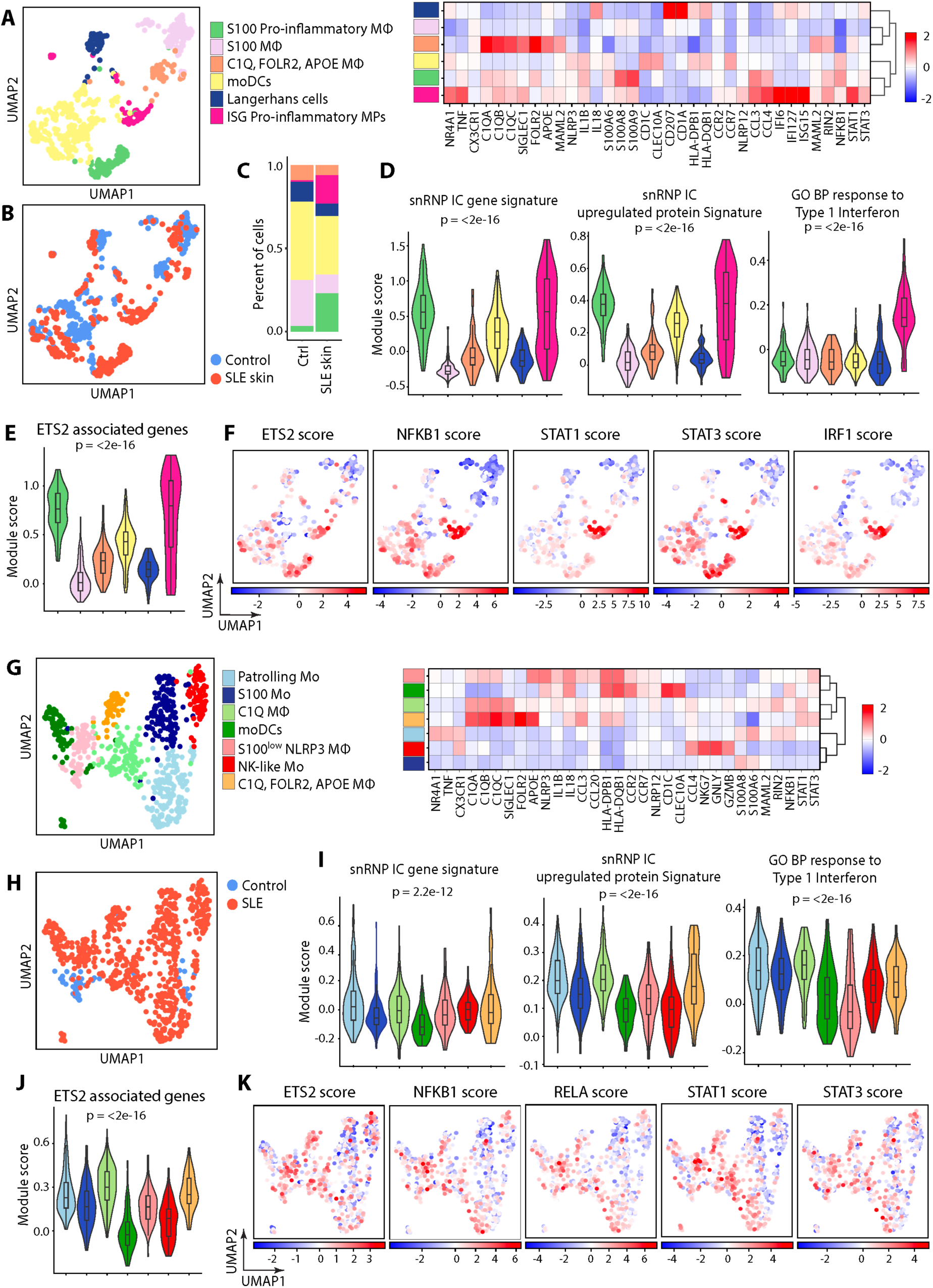
Single-cell transcriptomic profiling of mononuclear phagocytic cells (MPs) in acute cutaneous lupus and lupus nephritis identifies pro-inflammatory subsets with snRNP immune complex (IC) signature. Publicly available scRNA-seq data from acute cutaneous lupus (GSE186476) (A-F) and lupus nephritis for SLE (ImmPort, access code SDY997) (G-K) were analyzed. (A) UMAP visualization of MP clusters assigned to 6 subsets and scaled heatmap of mean expression of selected DEGs and markers used for cluster annotation. (B) UMAP colored by condition (blue = normal control skin, red = cutaneous lupus). (C) Stacked bar plot displaying the relative proportions of MP clusters across the two conditions. (D-E) Violin plots of module scores for expression of (D) snRNP IC gene signature, genes corresponding to upregulated differentially expressed proteins in response to snRNP IC stimulation, GO BP response to Type I Interferon gene set, and (E) ETS2 associated genes. (F) Feature plots illustrating inferred ETS2, NFKB1, STAT1, STAT3, and IRF1 transcription factor activities. (G) UMAP visualization of MP clusters in lupus nephritis assigned to 7 subsets and scaled heatmap of mean expression of select DEGs and markers used for cluster annotation. (H) UMAP colored by condition (blue = normal control kidney, red = lupus nephritis kidney). (I-J) Violin plots of module scores for expression of (I) the snRNP IC gene signature, genes corresponding to upregulated DEPs in response to snRNP IC stimulation, GO BP response to Type I Interferon gene set and (J) ETS2 associated genes. (K) Feature plots illustrating inferred ETS2, NFKB1, RELA, STAT1 and STAT3 transcription factor activities. Statistical comparisons of module scores were performed using the Kruskal-Wallis test.

TF activity inference identified increased activity of ETS2 and significant enrichment of ETS2 associated genes, which were derived from genes associated with ETS2 overexpression (*14*), in S100 pro-inflammatory MΦ and ISG pro-inflammatory MPs (Fig. 4E-F). While the activities of STAT1 and IRF1, a TF downstream of interferon receptors, were mainly increased in ISG pro-inflammatory MPs (Fig. 4F), increased activity of STAT3, and to a lesser extent NFKB1, was observed in both ISG pro-inflammatory MPs and S100 pro-inflammatory MΦ.

### MP subsets with lupus IC signature are found in lupus nephritis as determined by scRNA-seq analysis

We next explored MPs in lupus nephritis using publicly available scRNA-seq data from kidney biopsies from 24 patients with lupus nephritis (WHO class III and IV) and 10 control samples (living donor kidney biopsies) (*18*). MPs (n = 687 cells) were identified based on expression of *LYZ, CD14,* and *FCGR3A*. Clustering of these cells yielded seven clusters (Fig. 4G) which were assigned to the following cell types mainly based on DEGs (Fig. 4G heatmap): patrolling Mo, NK-like Mo, S100 Mo, C1Q MΦ, C1Q/FOLR2/APOE MΦ, S100^low^ NLRP3 MΦ, and moDCs. Of note, only a small number of these cells (n = 54) were from control kidney tissue (Fig. 4H). Expression of the snRNP IC gene signature was relatively modest with some subsets such as patrolling Mo, C1Q MΦ, NK-like Mo, and C1Q/FOLR2/APOE MΦ demonstrating higher expression compared to others (Fig. 4I). Similar findings were observed for the expression of dsDNA and Ro IC gene signatures from our microarray analysis (Fig. S4). Relatively higher expression of the snRNP IC protein signature was found in five subsets including patrolling Mo, S100 Mo, C1Q MΦ, C1Q/FOLR2/APOE MΦ, and S100^low^ NLRP3 MΦ (Fig. 4I). Patrolling Mo, S100 Mo, and C1Q MΦ demonstrated the highest expression of type I interferon inducible genes (Fig. 4I). Apart from S100 Mo, most subsets expressed additional inflammation-associated genes. For instance, S100^low^ NLRP3 MΦ demonstrated relatively high expression of *NLRP3*, *IL1B*, *IL18*, *CCL3*, and *CCL20* (Fig. 4G heatmap), whereas C1Q MΦ and C1Q/*FOLR2/APOE* MΦ showed relatively modest to high expression of these genes. Patrolling Mo demonstrated the highest expression of *NR4A1*, a gene characteristic of this subset (*10*), T*NF*, and *CX3CR1*. These four subsets demonstrated relatively high expression of ETS2-associated genes (Fig. 4J) as well as increased TF activity of ETS2, NFKB1, STAT1, and STAT3 (Fig. 4K), similar to Mo activated with lupus IC.

### scRNA-seq analysis of circulating Mo from lupus patients identifies a pro-inflammatory Mo subset with lupus IC signature

We explored whether similar pro-inflammatory Mo activated by lupus IC could be detected among circulating Mo from lupus patients using publicly available scRNA-seq data of peripheral blood mononuclear cells (PBMCs) from 162 lupus patients and 99 healthy controls (GSE174188) (*19*). A total of 240,115 Mo were identified based on the expression of *LYZ, CD14,* and *FCGR3A*. Clustering of these Mo yielded nine distinct clusters that were assigned to the following subsets primarily based on DEGs (Fig. 5A-B): C1Q/APOE Mo, patrolling Mo, NK-like Mo, EGR1/NLRP7 Mo, moDCs, pro-inflammatory Mo, S100 Mo, ISG Mo, and FOLR2/ISG Mo. Pro-inflammatory Mo demonstrated highest expression of inflammatory genes including *NLRP3*, *IL1B*, *IL6*, and *CCL3* (Fig. 5B). Lupus patients exhibited expansion of pro-inflammatory, ISG, and FOLR2/ISG Mo subsets compared to healthy controls, whereas S100 and EGR1/NLRP7 Mo subsets were relatively contracted in lupus patients (Fig. 5C). Pro-inflammatory Mo, predominantly found in lupus patients, demonstrated the highest expression of the snRNP IC gene and upregulated protein signatures derived from Mo stimulated with snRNP IC (Fig. 5D) as well as dsDNA and Ro IC gene signatures (Fig 5SA). Pro-inflammatory Mo also exhibited the highest expression of TLR signaling, NOD-like receptor signaling, and cytokine-cytokine receptor interaction KEGG pathway gene sets (Fig. S5B). Patrolling, C1Q/APOE, ISG and FOLR2/ISG Mo subsets exhibited the highest expression of type I IFN signature (Fig. 5D), whereas pro-inflammatory Mo had relatively lower expression compared to these Mo subsets, suggesting that lupus IC and type I IFN stimulation may independently affect Mo subsets in SLE. Similar to lupus IC-stimulated Mo, the pro-inflammatory Mo demonstrated increased inferred TF activity of NFKB and RELA (Fig. 5E) and highest expression of ETS2 associated genes (Fig. 5F). These findings implicate lupus IC in the activation and expansion of a subset of circulating Mo that expresses inflammatory genes, including *NLRP3*, *IL1B, IL18, CCL2, CCL3,* and *CCL4* (Fig 5B).

**Figure 5.**
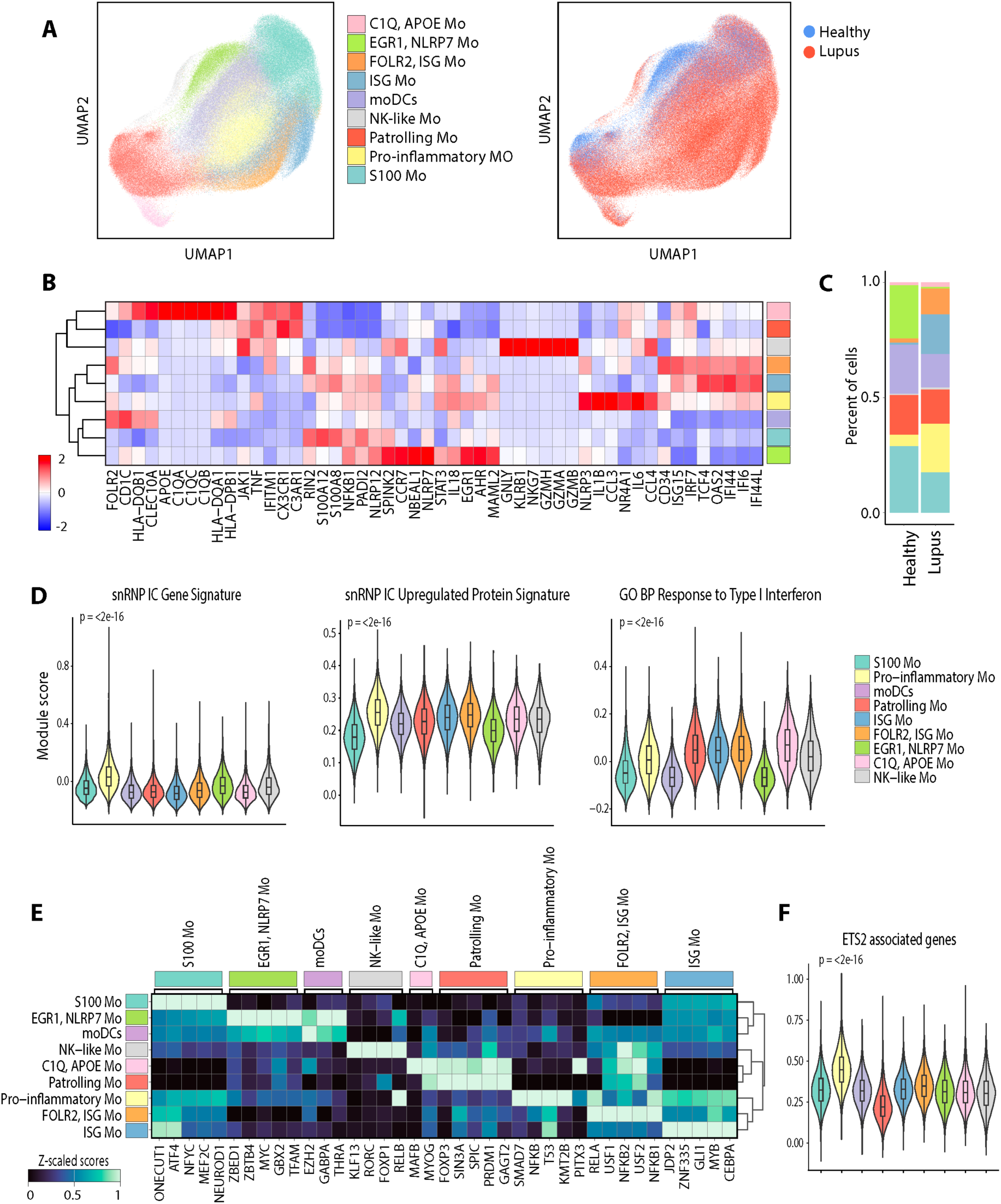
Single-cell transcriptomic profiling of circulating monocytes (Mo) in systemic lupus erythematous (SLE) identifies a pro-inflammatory subset with snRNP immune complex (IC) signature. Publicly available scRNA-seq data from the PBMCs of patients with SLE (n = 162) and healthy controls (n = 99) (GSE174188) were analyzed. (A) UMAP visualization of Mo clusters assigned to 9 subsets and UMAP colored by condition (blue = healthy control, red = Lupus). (B) Scaled heatmap of mean expression of selected DEGs and markers used for cluster annotation. (C) Stacked bar plot displaying relative proportions of Mo clusters across the two conditions. (D) Violin plots of module scores for expression of the snRNP IC gene signature, genes corresponding to upregulated differentially expressed proteins in response to snRNP IC stimulation, and GO BP response to Type I Interferon gene set. (E) Scaled heatmap of the top inferred transcription factor activities per Mo cluster. (F) Violin plot of module scores for expression of ETS2 associated genes. Statistical comparisons of module scores were performed using the Kruskal-Wallis test.

### The presence of CD68^+^ MΦ expressing NLRP3 is associated with clinical outcomes in lupus nephritis

We previously identified upregulation of NLRP3 at the gene and protein levels in bulk Mo stimulated with lupus IC, as well as the presence of NLRP3 expressing CD14^+^ cells in acute cutaneous lupus (*8*). In line with this finding, our scRNA-seq analyses revealed an expansion of pro-inflammatory Mo expressing *NLRP3* in Mo stimulated with lupus IC and in circulating Mo from lupus patients, as well as the presence of *NLRP3* expressing MPs in lupus nephritis. To extend these findings to the tissue protein level, we analyzed the relationship between MΦ expressing NLRP3 and clinical outcomes in lupus nephritis using our recently published IMC data consisting of kidney tissue samples from patients with lupus nephritis (WHO classes III and IV, n = 17) and normal control kidney tissues (n = 2) (*20*). In this dataset, lupus nephritis tissue samples were divided into treatment responder (urine protein/creatinine ratio < 0.5, normal serum creatinine (≤ 1.0 mg/dL), and prednisone ≤ 10 mg/day) and non-responder groups based on treatment outcomes (*20*). We detected NLRP3 expressing CD68^+^ MΦ in lupus nephritis tissues (Fig. 6A). Patients with lupus nephritis in the treatment non-responder group had higher levels of CD68^+^ NLRP3^+^ MΦ infiltration in the kidney compared to those in the treatment responder group (Fig. 6B), supporting a potential role for these cells in the pathogenesis of lupus nephritis and their association with clinical outcomes.

**Figure 6.**
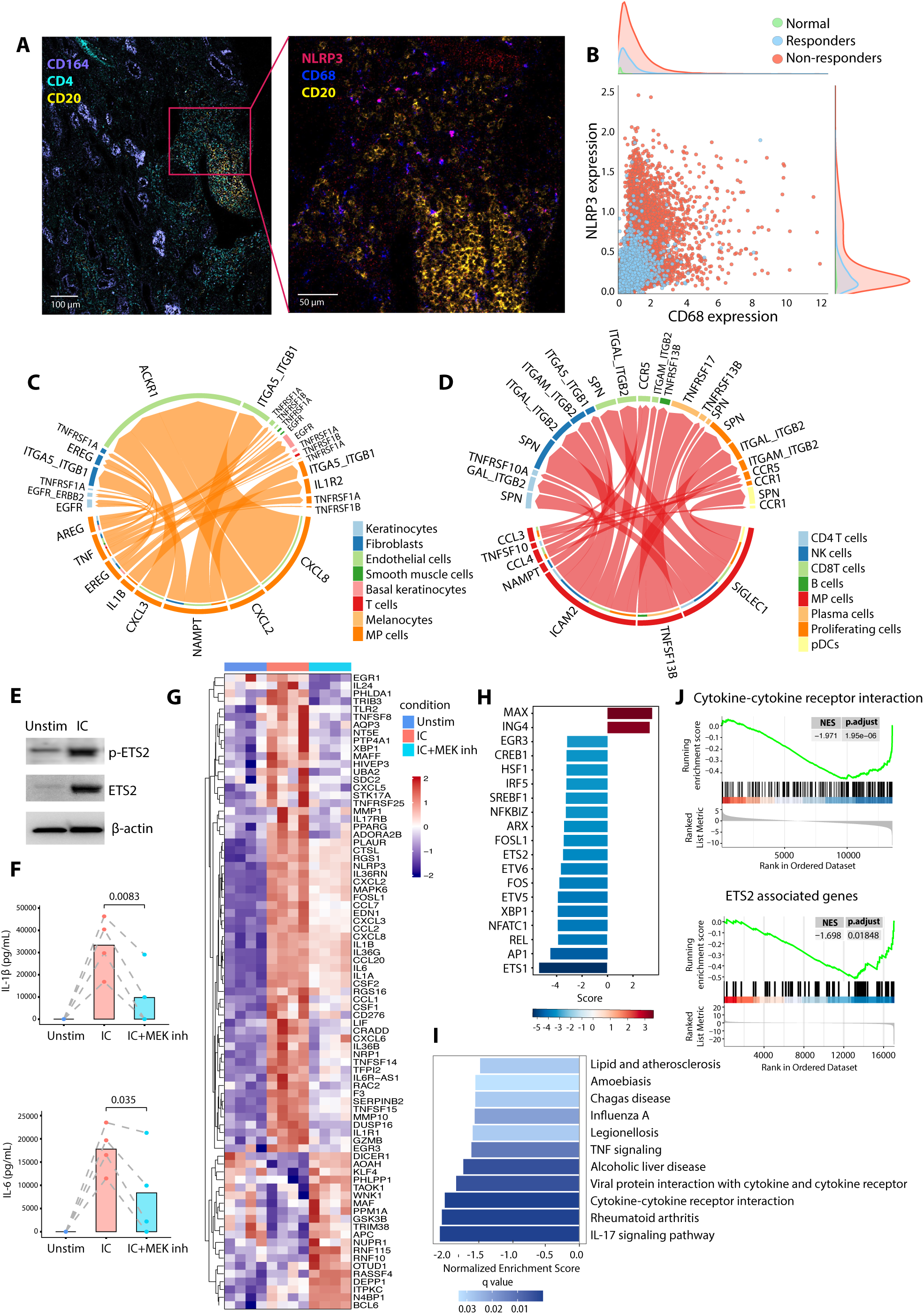
Pro-inflammatory NLRP3^+^ macrophages (MΦ) are associated with clinical outcomes in lupus nephritis and MEK inhibition attenuates lupus-IC induced activation of pro-inflammatory monocytes (Mo). (A) Imaging Mass Cytometry (IMC) image of renal biopsy from a treatment non-responder patient. Right panel shows a magnified view of the inset region in the left panel. (B) Bivariate scatter plot of CD45^+^ cells with marginal distributions illustrating the co-expression of CD68 (x-axis) and NLRP3 (y-axis) at the single-cell level, stratified by clinical outcome (7 treatment responders and 10 non-responders). (C-D) Cell to cell communication analysis of acute cutaneous lupus and lupus nephritis scRNA-seq data from Figure 4 using CellChat. (C) Chord diagram showing selected upregulated ligand-receptor interactions in cutaneous lupus (compared to control skin), with MPs as the source. (D) Chord diagram showing selected ligand-receptor interactions in lupus nephritis, with MPs as the source. (E) Western blot analysis of unstimulated and snRNP IC-stimulated Mo demonstrating ETS2 and phosphorylated ETS2 expression levels. Representative data from 2 independent experiments with 2 donors. (F) ELISA of IL-1β and IL-6 in the culture supernatants of unstimulated Mo (Unstim), snRNP IC-stimulated Mo (IC), and Mo pre-treated with the MEK inhibitor PD-0325901 prior to snRNP IC stimulation (IC + MEK inh). Numbers indicate *P* values obtained by the paired *t*-test. (G) Scaled heatmap of variance-stabilizing transformed (VST) counts showing expression of selected genes across Unstim, IC and IC + MEK inh Mo. (H) Bar plot showing inferred transcription factor activity from RNA-seq data comparing IC + MEK inh Mo vs IC Mo. (I) Bar plot showing negatively enriched KEGG pathways with MEK inhibition, identified by Gene Set Enrichment Analysis (GSEA). (J) GSEA plots for cytokine-cytokine receptor interaction and ETS2-associated genes gene sets.

### Lupus IC-stimulated MPs interact with immune and stromal cells in lupus skin and kidneys, and may be suppressed by IL-1 blockade

We explored whether molecules produced by lupus IC-stimulated Mo and MΦ could influence other immune and non-immune cells in lupus skin and kidney tissues by performing cell-cell communication analysis with CellChat (*21*), a tool that quantitatively infers intercellular communication networks from scRNA-seq data. In cutaneous lupus, communication between MPs and keratinocytes, fibroblasts, endothelial cells, smooth muscle cells, melanocytes, and T cells was upregulated via TNF, IL-1β, CXCL2 (growth-regulated oncogene beta or GRO-β), CXCL3 (GRO-γ), CXCL8 (IL-8), and nicotinamide phosphoribosyl transferase (NAMPT) (Fig. 6C and Fig. S6). These cytokines and chemokines were upregulated in Mo stimulated with snRNP IC, as determined by our transcriptomic and proteomic analyses. Similarly, lupus nephritis exhibited interactions between MPs and CD4^+^ T, CD8^+^ T, NK, B, plasma, and plasmacytoid DC (pDCs) cells via TNFSF13B (BAFF), TNFSF10 (TNF-related apoptosis inducing ligand or TRAIL), SIGLEC1, CCL3, CCL4, NAMPT, and ICAM2 (Fig. 6D and Fig. S7), which were upregulated in Mo stimulated with lupus IC.

We performed drug-target prediction analysis on the scRNA-seq datasets used in our study using Drug2Cell, a tool that characterizes drug-target expression at the single-cell level (*22*), to identify potential drug candidates that target pro-inflammatory Mo and MΦ. Mycophenolate mofetil, a cytotoxic drug commonly used in lupus, especially for deep organ involvement such as nephritis, was predicted to have minimal to modest targeting of S100 pro-inflammatory Mo and S100^low^ pro-inflammatory Mo (Fig. S8A), which expanded in lupus IC-stimulated Mo. Notably, this analysis identified IL-1 inhibitors, including canakinumab, an anti-IL-1β neutralizing monoclonal Ab, and rilonacept, a dimeric fusion protein that functions as a soluble decoy receptor for IL-1, as potential candidates for targeting these pro-inflammatory Mo subsets (Fig. S8A). Tacrolimus, a calcineurin inhibitor, was predicted to exert a non-selective effect on lupus IC-stimulated Mo. In lupus skin lesions, canakinumab and rilonacept were predicted to target S100 proinflammatory MΦ and ISG proinflammatory MPs (Fig. S8B), two expanded subsets (Fig. 4C), whereas the effects of these drugs on MP subsets in lupus nephritis were predicted to be less selective (Fig. S8C). In circulating Mo from lupus patients, the pattern of predicted drug targeting was similar to that observed in Mo stimulated with lupus IC, with selective targeting of pro-inflammatory Mo by canakinumab and rilonacept (Fig. S8D). As in lupus IC-stimulated Mo, the effect of mycophenolate on MPs in lupus skin, kidneys, and PBMCs was predicted to be minimal to modest. Similarly, tacrolimus was predicted to have substantial effects on multiple MP subsets, including both inflammatory and non-inflammatory subsets, in lupus nephritis and circulating Mo from lupus patients (Fig. S8C-D).

### Targeting ETS2 activation with MEK inhibition attenuates lupus IC-driven pro-inflammatory activation of Mo

Analysis of scRNA-seq data from both lupus-IC stimulated Mo and from MPs across different tissues from lupus patients implicated the transcription factor ETS2 as a master regulator of pro-inflammatory MP subsets. Of note, ETS2 was recently identified as a central regulator of human inflammatory MΦ (*14*). Western blot analysis demonstrated increased expression of ETS2 and phosphorylated (activated) ETS2 in Mo stimulated with snRNP IC (Fig. 6E). The MAPK/ERK Kinase (MEK) pathway mediates ETS2 activation by inducing phosphorylation (*23*). In a published study, the MEK inhibitor PD-0325901 substantially reduced ETS2-regulated genes in human inflammatory MΦ differentiated from Mo in the presence of GM-CSF, TNF-α, PGE2, and Pam3CSK4 (*14*). In line with these findings, PD-0325901 decreased IL-1β and IL-6 production from Mo stimulated with lupus IC (Fig. 6F). As determined by RNA-seq analysis, the MEK inhibitor PD-0325901 affected the expression of multiple inflammatory genes, including those encoding cytokines, chemokines, and *NLRP3* in snRNP IC-stimulated Mo (Fig. 6G). MEK inhibitor-treated cells also exhibited increased expression of genes encoding negative regulators of inflammation such as acyloxyacyl hydrolase (*AOAH*) (*24*), OTU deubiquitinase 1 (*OTUD1*) (*25*), and NEDD4 binding protein 1 (*N4BP1*) (*26*) (Fig. 6G). Inferred transcription factor activity analysis showed decreased activity of ETS2, ETS1, other downstream targets of MAP kinase signaling (AP1, FOS, FOSL1, CREB1) as well as NF-κB family associated transcription factors, including REL and NFKBIZ, with MEK inhibition (Fig. 6H). Additionally, inferred activity of ING4 (*27, 28*), an anti-inflammatory regulator that inhibits NF-κB signaling, was increased with MEK inhibition. GSEA demonstrated negative enrichment of many of the KEGG pathways that were enriched in lupus IC-activated pro-inflammatory Mo including “cytokine-cytokine receptor interaction”, “IL-17 signaling”, and “TNF signaling” (Figs. 6I-J) as well as ETS2 associated genes (Fig. 6J). Targeting ETS2 activation with MEK inhibition attenuated lupus IC-driven activation of pro-inflammatory Mo and may promote reprogramming to an anti-inflammatory state.

## Discussion

IC consisting of autoantibody and autoantigen likely play a critical role in the pathogenesis of lupus through activation of MPs, including Mo and MΦ, which can produce an array of inflammatory molecules. However, the global inflammatory changes induced in MPs activated by lupus IC, and whether such alterations are present in organ systems affected by lupus, have remained largely unknown. To address this, we have characterized transcriptomic and proteomic changes in Mo in response to lupus snRNP IC by analysis of scRNA-seq, gene expression microarray, bulk RNA-seq and proteomic data (Graphical Abstract). We further interrogated the skin, kidneys, and PBMCs of lupus patients for evidence of these changes *in vivo*. Our study identified substantial expansion of a Mo subset, referred to as S100 pro-inflammatory Mo, characterized by expression of inflammatory genes encoding cytokines, chemokines, and other inflammatory molecules such as NLRP3 in Mo stimulated with snRNP IC. The production of these inflammatory molecules by snRNP IC stimulated Mo was further validated at the protein level through in-depth proteomic analysis. Stimulation of Mo with other types of lupus-associated IC, comprised of dsDNA or Ro60, elicited upregulation of similar pro-inflammatory genes, highlighting the activation of pro-inflammatory Mo by different types of IC in lupus. Pathway and gene set enrichment analyses combined with TF activity inference identified TLR, NOD-like receptor, and JAK-STAT signaling as main pathways, and NF-kB and ETS2 as master regulator TFs, involved in mediating the inflammatory changes in lupus IC-activated Mo. Analysis of scRNA-seq datasets revealed the presence and expansion of pro-inflammatory Mo and MΦ subsets with similar inflammatory gene expression profiles across the skin, kidneys, and PBMCs of lupus patients, suggesting a shared lupus IC-driven mechanism underlying this phenomenon. Overall, our findings corroborate the role of lupus IC in the induction and expansion of highly inflammatory Mo and MΦ subsets across multiple affected organs in lupus, providing a scientific rationale for therapeutically targeting these cells.

In our scRNA-seq analysis of human Mo stimulated with snRNP IC, we observed a marked expansion of pro-inflammatory Mo expressing S100 genes, including *S100A8* and *S100A9*, from approximately 2% to over 50% in response to snRNP IC. In contrast, a Mo subset expressing the same S100 genes as well as *NLRP12* but lacking expression of inflammatory genes contracted to a similar extent, suggesting a possible reciprocal relationship between these Mo subsets. This notion was supported by pseudotime trajectory analysis, which showed differentiation of S100 Mo without expression of inflammatory genes, the most abundant subset in unstimulated Mo, into S100 pro-inflammatory Mo upon activation with lupus IC, while maintaining expression of *S100A8* and *S100A9*. The transformation into pro-inflammatory Mo was accompanied by downregulation of *NLRP12*, a negative regulator of Mo activation and cytokine production (*12*), suggesting that NLRP12 plays a role in maintaining S100 Mo in a quiescent state in the absence of IC stimulation. We observed expansion of pro-inflammatory MΦ and Mo subsets, including those expressing S100, in the skin and peripheral blood of lupus patients, supporting the biological and clinical significance of our *ex vivo* Mo stimulation findings. *S100A8* and *S100A9* encode calgranulins A and B which together form calprotectin, a heterodimer with antimicrobial and pro-inflammatory properties (*29*). Calprotectin serves as a ligand for pattern recognition receptors (PRRs) including the receptor for advanced glycation endproducts (RAGE), TLR4, and CD33 (*30*). Extracellular S100A8 and S100A9 can activate the NF-κB pathway via TLR4, resulting in the production of pro-inflammatory cytokines (*31*). Increased serum levels of S100A8 and S100A9 were found in lupus patients with anti-dsDNA Abs and nephritis (*32*). A recent urine proteomics study identified strong correlations between urinary S100A9 and MCP-1 (CCL2), a chemokine highly produced by snRNP IC-stimulated Mo, and the presence and severity of glomerular lesions and interstitial inflammation in lupus nephritis (*33*). Together, these findings suggest that S100 Mo serve a surveillance function at rest and when activated, transform into a pro-inflammatory phenotype contributing to tissue inflammation and immune cell recruitment via secretion of cytokines, chemokines, and other inflammatory mediators.

We identified populations of pro-inflammatory Mo and MΦ in the skin, kidneys, and PBMCs of lupus patients, which expressed the snRNP IC gene and protein signatures. Despite some similarities, MPs also exhibited organ-specific cellular heterogeneity, which could be related to several possible contributing factors including acuity vs. chronicity of stimulation, the presence or absence of additional stimuli such as type I interferons, and the influence of tissue-specific immune and non-immune microenvironments. Of note, some subsets of expanded MPs, including S100 pro-inflammatory MΦ and pro-inflammatory Mo in the skin and PBMCs of lupus patients, respectively, demonstrated increased expression of the snRNP IC gene signature though low expression of type I interferon inducible genes. These findings suggest that lupus IC and type I IFN stimulation may independently affect Mo subsets, highlighting the potential need for targeting lupus IC-mediated Mo activation in addition to blocking type I IFN signaling in SLE.

Our gene set, biological pathway, and TF activity inference analyses implicated several pathways, including NF-κB, TLR, NOD-like receptor, and TNF signaling pathways, in the activation of pro-inflammatory Mo and MΦ by lupus IC. Notably, these pathways were enriched in Mo activated by dsDNA and Ro IC as well. These findings are consistent with our published studies demonstrating the role of TLR7/8/9 and NF-κB in driving activation of human Mo in response to lupus IC (6–8). Beyond mediating the production of cytokines, chemokines, and growth factors, the NF-κB pathway also promotes the differentiation, activation and cell survival of Mo and MΦ (*34*). Of note, ETS2, recently identified as a critical master regulator of human inflammatory MΦ and implicated in immune-mediated diseases (*14*), may also play a role in SLE. In our study, S100 pro-inflammatory Mo activated by lupus IC as well as populations of pro-inflammatory Mo and MΦ identified in the skin and kidneys of lupus patients exhibited increased expression of ETS2-associated genes, which include *IL1B, IL6*, *TNF, S100A8, S100A9, MMP9, NFKB1,* and *IRF1*. These findings were further supported by ETS2 transcription factor activity inference analysis. Indeed, we observed increased protein expression of both ETS2 and phosphorylated ETS2 in Mo stimulated with lupus IC. Currently, direct inhibitors of ETS2 are not available. As ETS2 phosphorylation and activation depend on MAP kinase signaling (*23*), we targeted an upstream regulator of ETS2 using a MEK inhibitor. MEK inhibition not only attenuated lupus IC-driven expression of pro-inflammatory genes in Mo but may also promote reprogramming toward an anti-inflammatory state.

Our spatial analysis of lupus nephritis using IMC showed that kidney biopsies from patients who did not respond to treatment exhibited greater infiltration of NLRP3^+^CD68^+^ MΦ compared to biopsies from treatment-responsive patients, suggesting a potential role for these cells in the pathogenesis and clinical outcomes of lupus nephritis. We previously demonstrated the role of lupus IC in activating the NLRP3 inflammasome in Mo, leading to IL-1β and IL-18 production (*6, 7*). These cytokines can further promote the activation of innate and adaptive immune cells. For instance, IL-1β can enhance the proliferation, differentiation, migration, and effector functions of CD4^+^ and CD8^+^ T cells (*35*), while IL-18 activates B cells and induces self-reactive IgM and IgG Ab responses (*36*). Notably, serum IL-18 levels have been associated with active renal disease and irreversible organ damage in patients with SLE (*37*). Our drug prediction analysis identified IL-1 inhibitors as potential candidates to abrogate IC-mediated activation of MPs in SLE. These findings support the rationale of targeting NLRP3 in lupus, which is further supported by studies linking *NLRP3* gene polymorphisms to genetic susceptibility of SLE and NLRP3 levels to tissue damage in lupus nephritis (*38*). Studies in lupus prone mice also demonstrated improvement of proteinuria, renal histologic lesions, and podocyte effacement with inhibition of NLRP3 (*39*). Moreover, clinical trials of NLRP3 inflammasome inhibition in various inflammatory diseases have shown encouraging results (*40*). While NLRP3 inhibitors represent a promising therapeutic option, targeting the broader pro-inflammatory transcriptomic and proteomic changes in MPs induced by lupus IC may offer further alleviation of tissue inflammation and damage in SLE.

We acknowledge limitations of our transcriptomic analysis, which need to be complemented by additional protein assays to better understand the biological significance of our findings. Indeed, we demonstrated a significant correlation between DEGs and DEPs in Mo stimulated with snRNP IC based on gene expression microarray and high-plex proteomic (i.e., SomaScanR® assay) analyses. Since the proteomic assay was performed on culture supernatants from stimulated Mo, intracellular protein analysis could provide additional insight into changes in non-secreted intracellular molecules, such as signaling molecules, in Mo in response to lupus IC.

Taken together, our study identified lupus IC-driven pro-inflammatory responses at both the transcriptomic and proteomic levels in human Mo, along with altered cellular heterogeneity following activation by lupus IC. We observed expansion of distinct Mo and MΦ populations bearing lupus IC-associated gene and protein signatures, which encompass key inflammatory molecules such as NLRP3, IL-1β, and IL-18, in the skin, kidneys, and peripheral blood of lupus patients. The transcription factor ETS2 may act as a master regulator of these pro-inflammatory Mo and MΦ populations. The clinical significance of these cell populations in lupus nephritis is further supported by the association between increased infiltration of NLRP3^+^CD68^+^ MΦ and poor treatment response. Collectively, these findings support a central role for lupus IC in driving the induction and expansion of highly inflammatory Mo and MΦ subsets across multiple affected organs in SLE, providing scientific rationale for therapeutically targeting these cells.

## Materials and Methods Summary

Detailed materials and methods are provided in the Supplementary Materials.

### Human subjects and Ethics

This work was approved by the institutional review boards (IRB) of Yale University and the Kyungpook National University Hospital and conducted in accordance with the principles of the Declaration of Helsinki. For scRNA-seq, gene expression microarray, and proteomics (SomaScan® assay, SomaLogic Operating Co, Inc., Boulder, CO) analyses, peripheral blood was obtained from healthy adult subjects (6–8). Anti-U1-snRNP Ab-positive sera were obtained from the peripheral blood of patients with SLE who met the 2019 American College of Rheumatology/European League Against Rheumatism Classification Criteria for SLE (*41*) or from the L2 Diagnostic Laboratory. Written informed consent was obtained from all human subjects.

### Mo stimulation for scRNA sequencing, gene expression microarray, SomaScan® assay, multiplex immunoassay, and Western blot

Untouched Mo (CD14^+^CD16^-^) were isolated from the peripheral blood of healthy adult subjects using a kit (Stemcell Technologies, Canada). Mo (1 x 10^5^) were resuspended in 200 μl of RPMI 1640 media supplemented with 10% FCS, penicillin, and streptomycin. For scRNA-seq and Western blot, Mo were incubated for 2 hours with or without a combination of U1-snRNP (5 μg/ml, AroTec Diagnostics Limited, New Zealand) and pooled anti-U1-snRNP Ab+ serum (final concentration of 2%) (referred to as snRNP immune complex or IC) (*6*). For Western blot, protein extracts were separated by SDS-PAGE and transferred onto PVDF membranes. Membranes were probed with primary antibodies against total ETS2 and phospho-ETS2 (Thr72) (Invitrogen), and β-actin (Santa Cruz Biotechnology) as a loading control, followed by staining with HRP-conjugated secondary antibodies (Santa Cruz Biotechnology) and visualization with Pierce ECL Western Blotting Substrate (Thermo Scientific). For multiplexed scRNA-seq, the incubated cells were stained for 30 minutes with cell hashtag Abs (Biolegend, San Diego, CA) conjugated to a unique barcode sequence and profiled using the 10x Genomics Chromium platform at the Yale Center for Genome Analysis (YCGA). The raw scRNA-seq data were processed and aligned to the human reference genome, hg38, using Cell Ranger (v7.1.0, 10x Genomics) (*42*). The filtered gene-cell barcode count matrices were further analyzed using Seurat (version 4.2.1) package (*43*) (see scRNA-seq data analysis below and Supplementary Materials and Methods).

For gene expression microarray analysis, Mo were incubated for 3 hours with or without snRNP IC, human genomic dsDNA (5 μg/ml) and pooled anti-dsDNA Ab+ serum (dsDNA IC), or Ro60 (5 μg/ml, AroTec Diagnostics Limited) and pooled anti-Ro60 Ab+ serum (Ro60 IC) (final concentration of 5 %). Anti-dsDNA Ab+ and Ro60 Ab+ sera were obtained from the L2 Diagnostic Laboratory. Total RNA was isolated from the stimulated cells using a Qiagen RNeasy mini kit (RNA integrity numbers∼10), amplified, and then hybridized to an Illumina HumanHT-12v4 Expression BeadChip (Illumina, Inc., San Diego, CA). Raw expression data were normalized using the quantile method (*44*), and batch effect was corrected using ComBat (*45*). Differential expression analysis was performed using the integrated hypothesis testing method (*46*). False discovery rate (FDR) was estimated by Storey’s method (*47*). DEGs were selected using thresholds of FDR < 0.05 and log2-fold-change ≥ 1. Three independent comparisons for each IC were performed and identified each set of DEGs. Functional enrichment analysis of the DEGs was performed using DAVID (*48*), and significantly enriched KEGG pathways were selected based on a threshold p-value < 0.01.

For the SomaScan® assay, culture supernatants of Mo incubated for 14 hours with or without snRNP IC were analyzed using a customized 1.45k SomaScan® assay panel (Supplementary Table). Protein levels were normalized using adaptive normalization by maximum likelihood, following SomaLogic’s standard protocol. The data were log-transformed, and baseline serum proteins were removed prior to differential protein expression analysis, which was conducted using the linear modeling approach implemented with limma (version 3.58.1) (*49*). *P*-values were adjusted for multiple testing with Benjamini-Hochberg correction (*50*).

For the multiplex immunoassay, supernatants from Mo incubated for 14 hours with and without snRNP IC were shipped to Eve Technologies (Calgary, Alberta, Canada) on dry ice. Cytokine and chemokine levels were measured in singlet using the Human Cytokine/Chemokine 96-Plex Discovery Assay® Panel (HD96). The heatmap of concentration levels was generated using ComplexHeatmap (version 2.18.0) (*51*).

### Inhibition of lupus-IC induced activation of Mo and bulk RNA sequencing

Human Mo freshly isolated from PBMCs were pretreated with the MEK inhibitor PD-0325901 (mirdametinib; MedChemExpress, LLC) for 30 minutes, followed by stimulation with snRNP ICs. For transcriptomic analysis, cells were stimulated for 4 hours, after which total RNA was extracted and used to generate cDNA and barcoded sequencing libraries with the TruSeq Stranded mRNA Kit (Illumina) according to the manufacturer’s instructions. Libraries were sequenced on an Illumina HiSeq 4000 platform using a 2 × 100 bp paired-end configuration. For cytokine analysis, Mo were stimulated for 12 hours, and levels of IL-1β and IL-6 in culture supernatants were measured by ELISA. Normality was assessed using the Shapiro-Wilk test. A paired *t*-test was used to compare cytokine levels in the presence and absence of MEK1/2 inhibitor.

Pseudoalignment-based transcript quantification of the raw bulk RNA sequencing data was performed using Salmon (v1.10.3) (*52*) with the *Homo sapiens* GRCm38 reference transcriptome.

The resulting transcript-level abundance estimates were imported into R (v4.5.1) and aggregated to gene-level counts using tximport (v1.38.2) (*53*). Differential expression analysis was conducted with DESeq2 (v1.50.2) (*54*) using the default Wald test. P-values were adjusted for multiple testing using the Benjamini-Hochberg method (*50*). Differentially expressed genes with adjusted p-value < 0.05 were considered significant. Heatmaps of gene expression were generated from variance-stabilizing transformed (VST) counts. Transcription factor activity inference was performed using the univariate linear modeling (ULM) method implemented with decoupler (v2.1.2) (*13*). Gene set enrichment analysis (GSEA) (*55*) was performed using clusterProfiler (v4.18.4) (*56*) with KEGG pathway gene sets. Gene sets with a false discovery rate (FDR) < 0.05 were considered significantly enriched.

### scRNA-seq data analysis

In addition to our own scRNA-seq data from above, publicly available scRNA-seq datasets were obtained and analyzed. scRNA-seq data from skin biopsies of patients with active cutaneous lupus and healthy normal skin were acquired from the National Center for Biotechnology Information (NCBI) Gene Expression Omnibus (GEO) data repository (GSE186476) (*17*). scRNA-seq data of kidney biopsies from patients with lupus nephritis and control kidney biopsy samples were acquired through the NIH Accelerated Medicine Program (AMP) for SLE (ImmPort, access code SDY997) (*18*). scRNA-seq data from PBMCs of patients with SLE and healthy controls were acquired from the NCBI GEO data repository (GSE174188) (*19*).

The filtered gene-cell barcode count matrices were analyzed using Seurat (version 4.2.1) pipelines including pre-processing, normalization, scaling, dimensionality reduction and clustering (*43*). Filtering, pre-processing, normalization, scaling, dimensionality reduction and clustering of these scRNA-seq datasets were performed as outlined in the Supplementary Materials and Methods. Biological processes and gene-gene correlations were assessed using Gene Set Enrichment Analysis (GSEA) (*55*). GSEA of Gene Ontology Biological Processes (GO BP) and Kyoto Encyclopedia of Genes and Genomes (KEGG) pathways was performed using clusterProfiler (version 4.2.2) (*56*). Trajectory analysis was conducted using scFates (version 1.0.6) following standard pipelines (*15*). Transcription factor activity was inferred using decoupleR (version 2.8.0) (*13*). Drug-target interactions were predicted using Drug2cell (version 0.1.1), which scores interactions based on expression of target genes (*22*). Additional plots and visualizations were generated using Scanpy (version 1.9.3) (*57*). Cell-cell communication was performed using CellChat (*21*).

### Imaging Mass Cytometry (IMC) analysis of kidney tissues

Pre-existing IMC data consisting of kidney biopsies from patients with lupus nephritis (WHO class III and class IV, n = 17) and normal control kidneys (n = 2) (*20*) were analyzed for the presence of cells expressing CD68 and NLRP3. Subject demographics, relevant clinical laboratory values, biopsy findings, and treatment regimens for the treatment responders (n=7) and non-responders (n=10) have been previously described by Lee *et al*. (*20*). Processing, cell detection, cell segmentation, cell measurements, cell classification and identification of the CD45+ cell subset were performed as outlined by Lee *et al.* (*20*). The CD45+ immune cell subset was interrogated for expression of CD68 and NLRP3, which was visualized using a bivariate scatter plot generated with the Python-based Seaborn (version 0.13.2) package (*58*).

## Statistical analysis

Details on statistical analyses were provided in Supplementary Materials and Methods. Statistical analyses were performed using R (version 4.5.1) and Python (version 3.11.14). For scRNA-seq, differential gene expression analysis was performed using the Wilcoxon rank-sum test, with p-values adjusted for multiple testing using Bonferroni correction (*59*). Comparisons of gene set module scores across cell clusters in scRNA-seq data were performed using the Kruskal-Wallis testing functionality of ggpubr (version 0.6.0) (*60*). For differential protein expression analysis, *P*-values were adjusted for multiple testing with Benjamini-Hochberg correction(*50*). Correlation analyses were conducted using Spearman’s rank correlation. Statistical significance was defined as *P*-values < 0.05.

## Data Availability

All data are available in the main text or the supplementary materials. Single-cell RNA-seq and bulk RNA-seq data will be publicly available through the NCBI Gene Expression Omnibus (GEO). Gene expression microarray data are publicly available through the NCBI GEO under accession code GSE315667. We also utilized the following publicly available datasets: the AMP dataset (ImmPort, access code SDY997), GSE186476, and GSE174188.

## List of Supplementary Materials

-Figures S1 to S8

-Table S1

-Supplementary Materials and Methods

## Author Contributions

LO and MSS designed the study, performed the experiments, analyzed and interpreted the results, and participated in writing the manuscript; HC, WJS, HK, JY,MK, WB, MF, JGA, HJP, JJS, JPY, ED, SU, JGA, JC, CL, MXD, FK, JLG, NK, RB and SY analyzed and interpreted the results, and participated in writing the manuscript; IK designed the study, analyzed and interpreted the results, participated in writing the manuscript and supervised the research. LO, MSS, and IK have verified the underlying data. All authors read and approved the final version of the manuscript.

