## Supplementary Figures for "Lupus Immune Complexes Drive Distinct Pro-Inflammatory Monocyte and Macrophage Populations Independent of Type I Interferon"

### Supplementary Figure 1

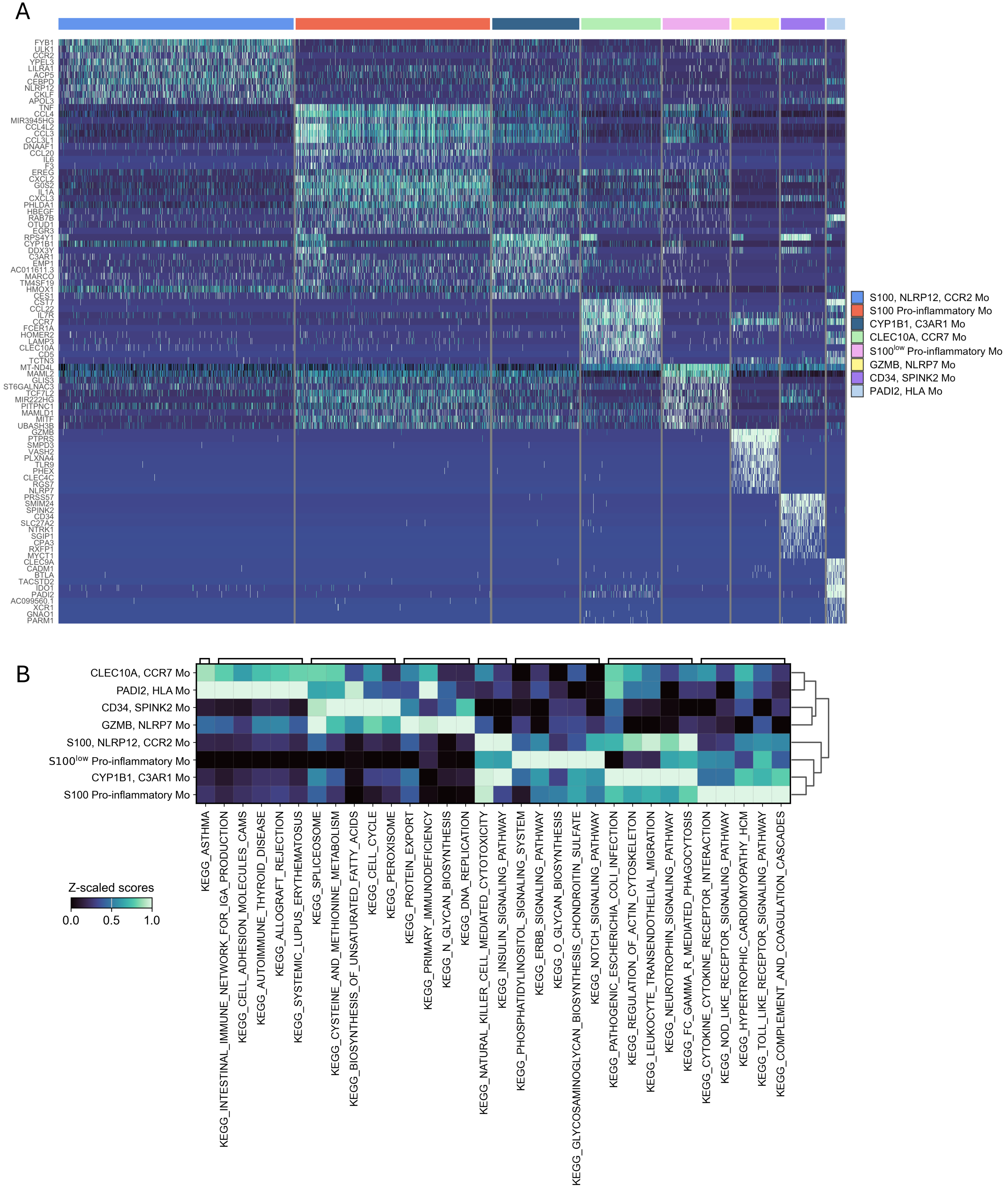


Supplementary Figure 1. Transcriptomic characterization of monocytes (Mo) stimulated with snRNP immune complex (IC) by single cell RNA-seq analysis. (A) Heatmap showing the top differentially expressed genes (DEGs) per Mo cluster. (B) Scaled heatmap of top enriched KEGG pathways per cluster

### Supplementary Figure 2

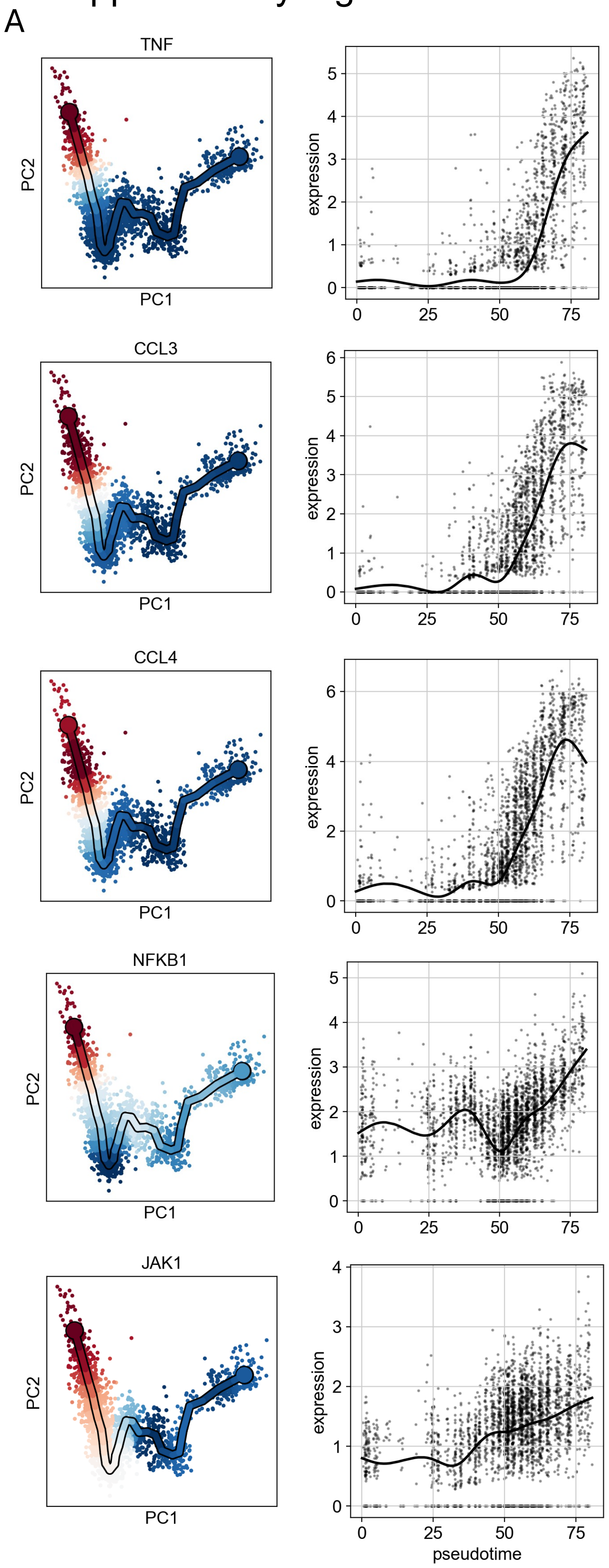


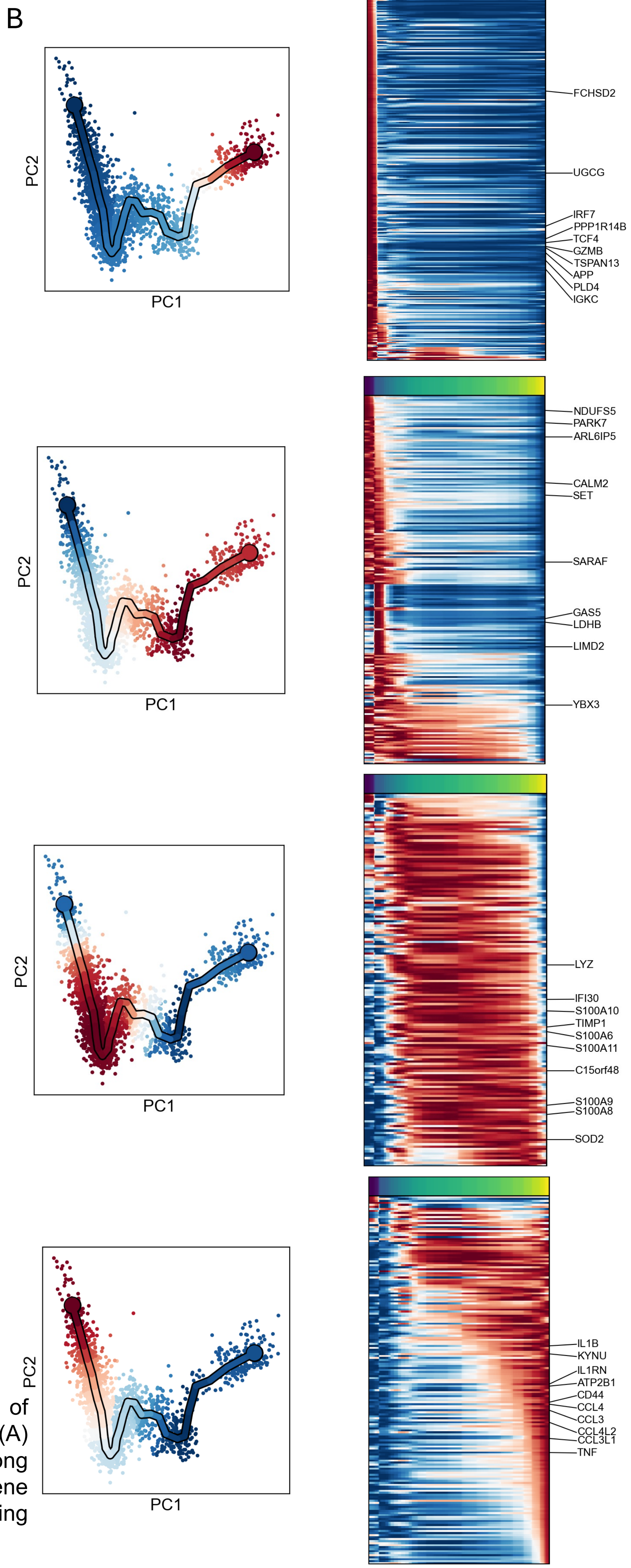


Supplementary Figure 2. Pseudotime trajectory analysis of snRNP IC stimulated monocytes analyzed by scRNA-seq. (A) Expression of *TNF*, *CCL3*, *CCL4*, *NFKB1*, and *JAK1* along pseudotime progression. (B) Heatmaps showing dynamic gene expression along pseudotime progression, with accompanying PCA plots indicating position along the trajectory.

### Supplementary Figure 3

A

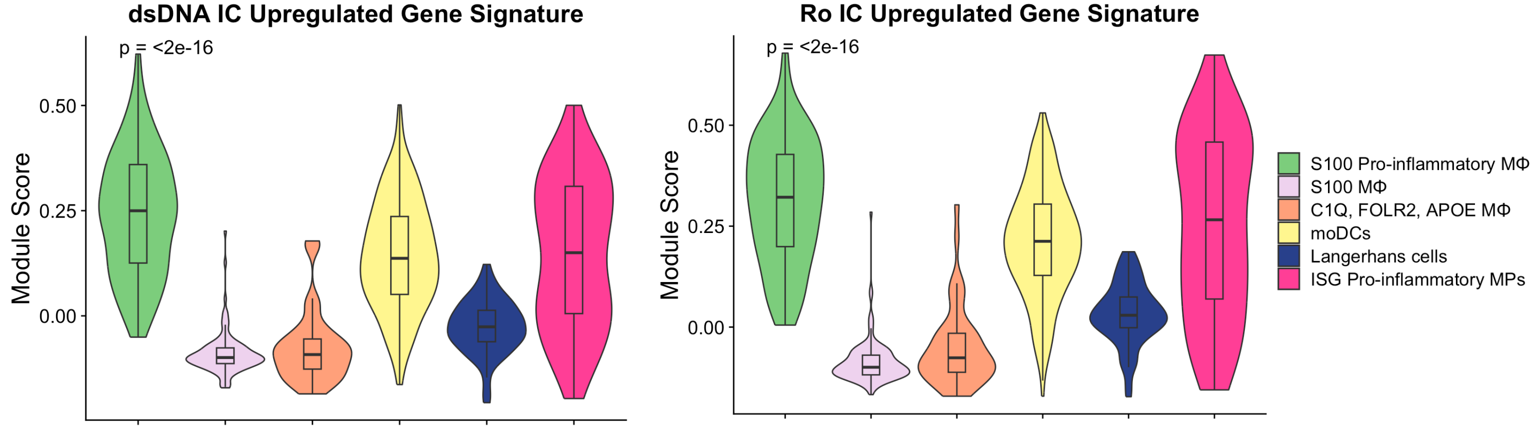


B

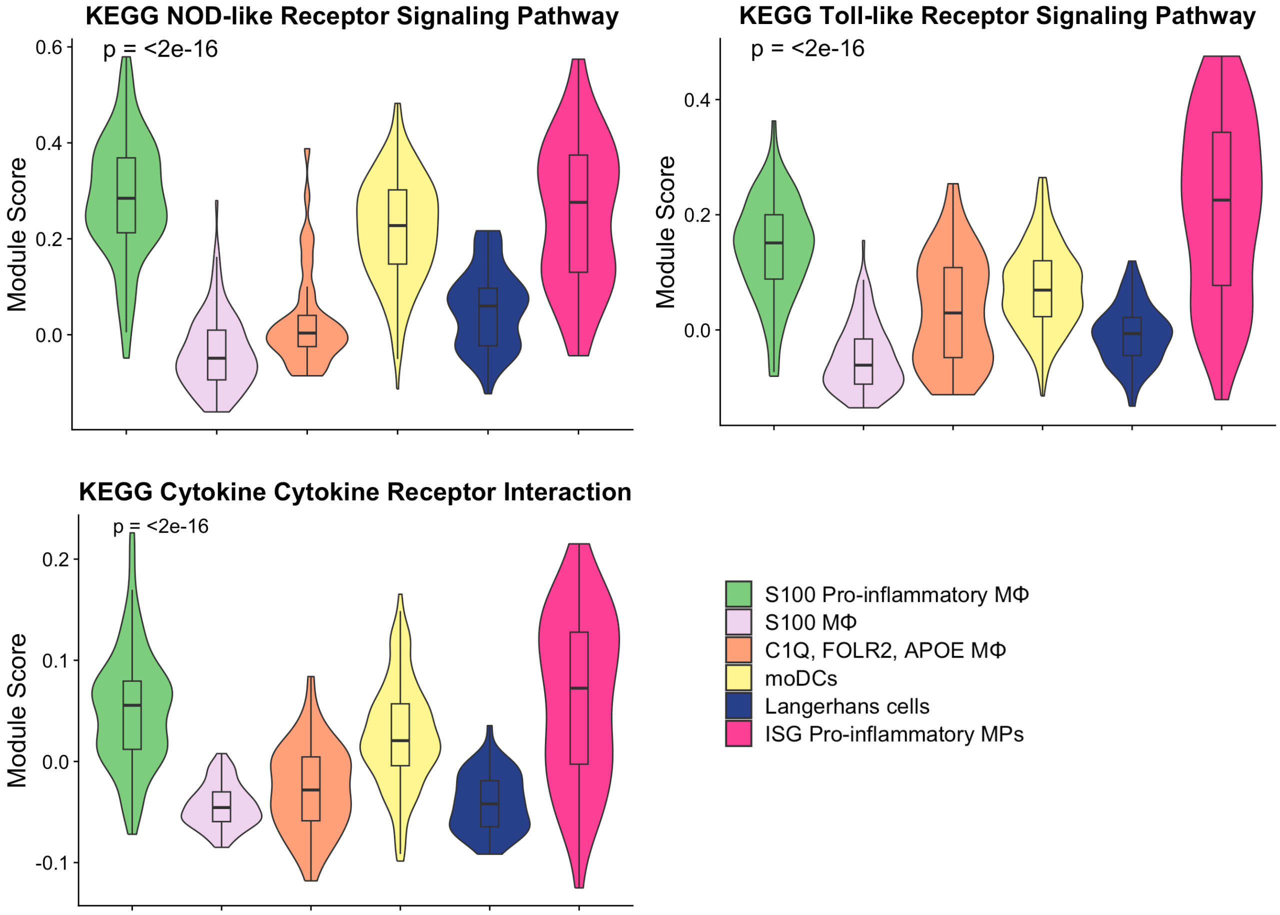


Supplementary Figure 3. Immune complex signature and pathway analysis of mononuclear phagocytic cells (MPs) in acute cutaneous lupus analyzed by scRNA-seq (data from GSE186476 - see details in Materials and Methods). (A) Violin plots of module scores for expression of upregulated genes identified from microarray analysis of Mo stimulated with dsDNA IC and Ro IC. (B) Violin plots of module scores for expression of KEGG NOD-like receptor signaling pathway, Toll-like receptor signaling pathway, and Cytokine-cytokine receptor interaction gene sets. Statistical comparisons of module scores were performed using the Kruskal-Wallis test.

### Supplementary Figure 4

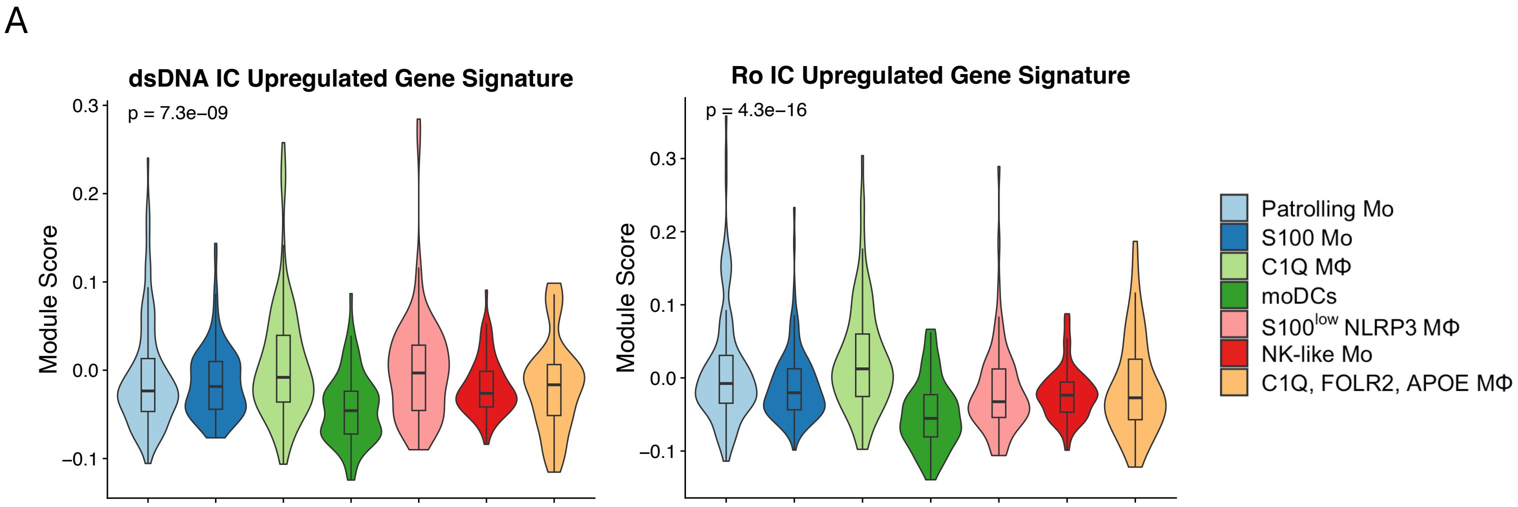


Supplementary Figure 4. Immune complex signatures among mononuclear phagocytic cells (MPs) in lupus nephritis analyzed by scRNA-seq (data from the NIH Accelerated Medicine Program (AMP) for SLE (ImmPort, access code SDY997)). (A) Violin plots of module scores for expression of upregulated genes identified from microarray analysis of Mo stimulated with dsDNA IC and Ro IC. Statistical comparisons of module scores were performed using the Kruskal-Wallis test.

### Supplementary Figure 5

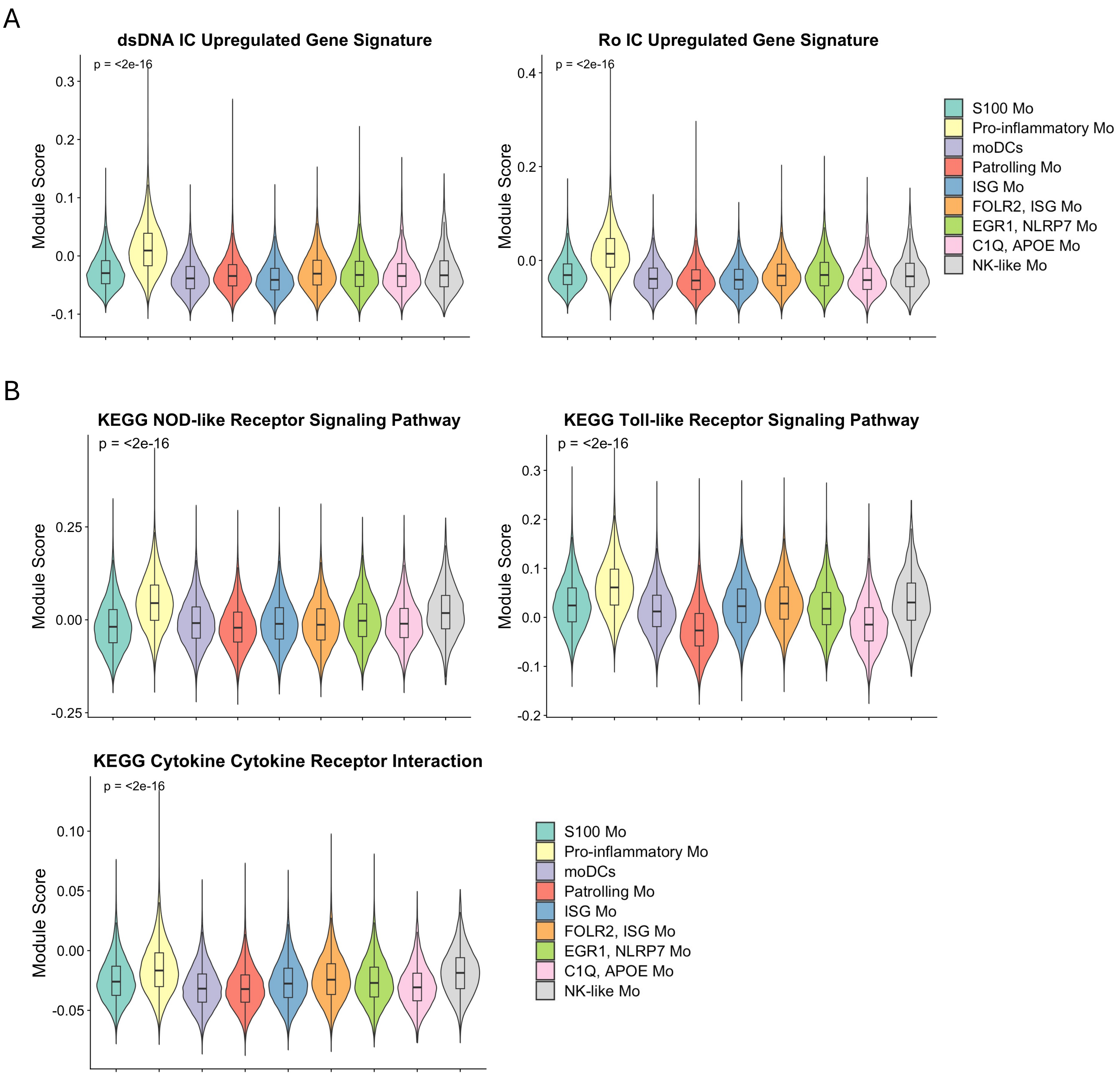


Supplementary Figure 5. Immune complex signature and pathway analysis of circulating monocytes (Mo) in Systemic Lupus Erythematous (SLE) analyzed by scRNA-seq (GSE174188, see details in Materials and Methods). (A) Violin plots of module scores for expression of upregulated genes identified from microarray analysis of Mo stimulated with dsDNA IC and Ro IC. (B) Violin plots of module scores for expression of KEGG NOD-like receptor signaling pathway, Toll-like receptor signaling pathway, and Cytokine-cytokine receptor interaction gene sets. Statistical comparisons of module scores were performed using the Kruskal-Wallis test.

### Supplementary Figure 6

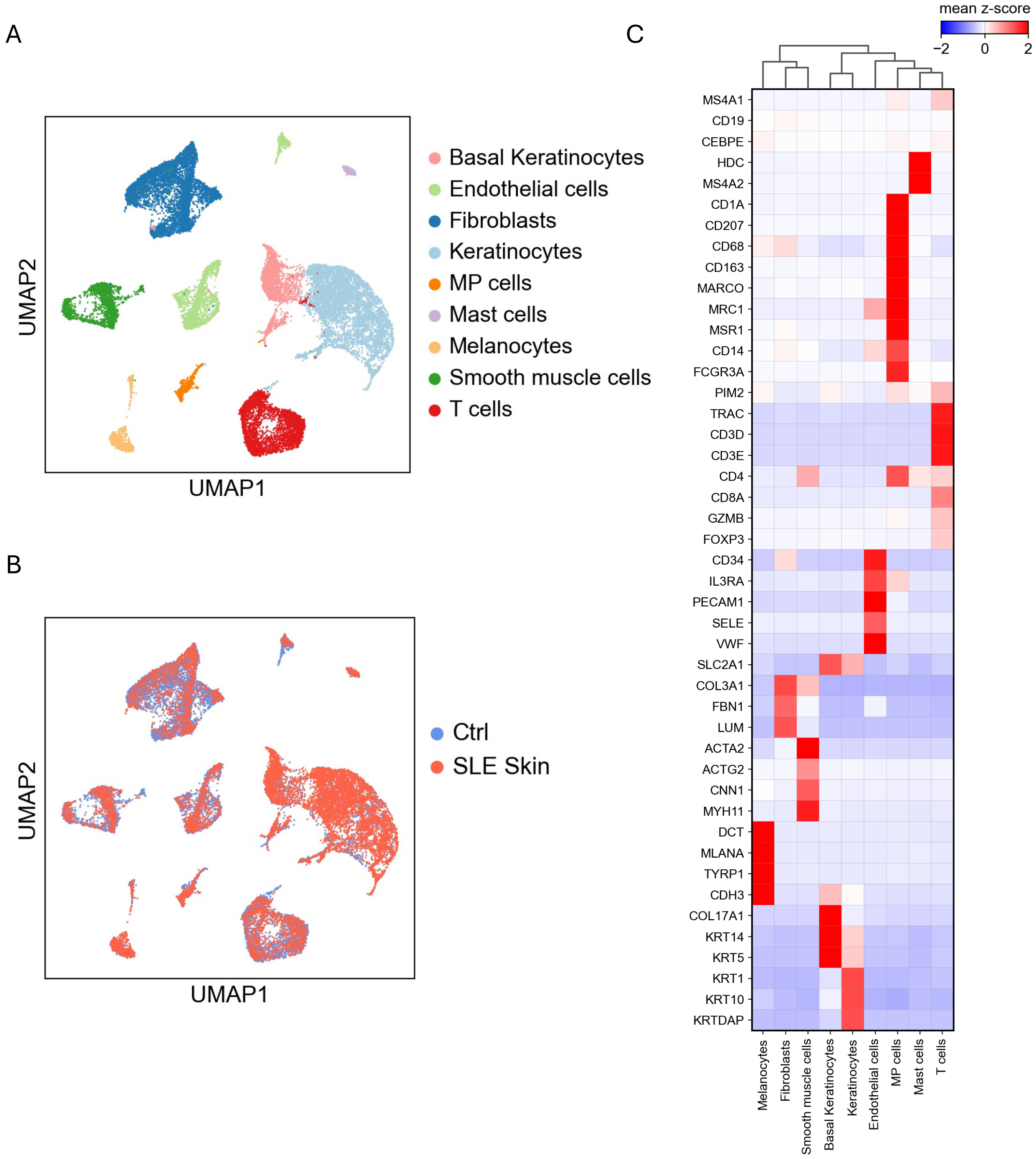


Supplementary figure 6. Single-cell profiling of acute cutaneous lupus by scRNA-seq (data from GSE186476 - see details in Materials and Methods). (A) UMAP visualization of cells assigned to 9 cell types based on their transcriptomic profiles (MP = Mononuclear Phagocytic). (B) UMAP colored by condition (blue = healthy control skin, red = cutaneous lupus). (C) Scaled heatmap of mean expression of select DEGs and markers used for cell type annotation.

Supplementary Figure 7

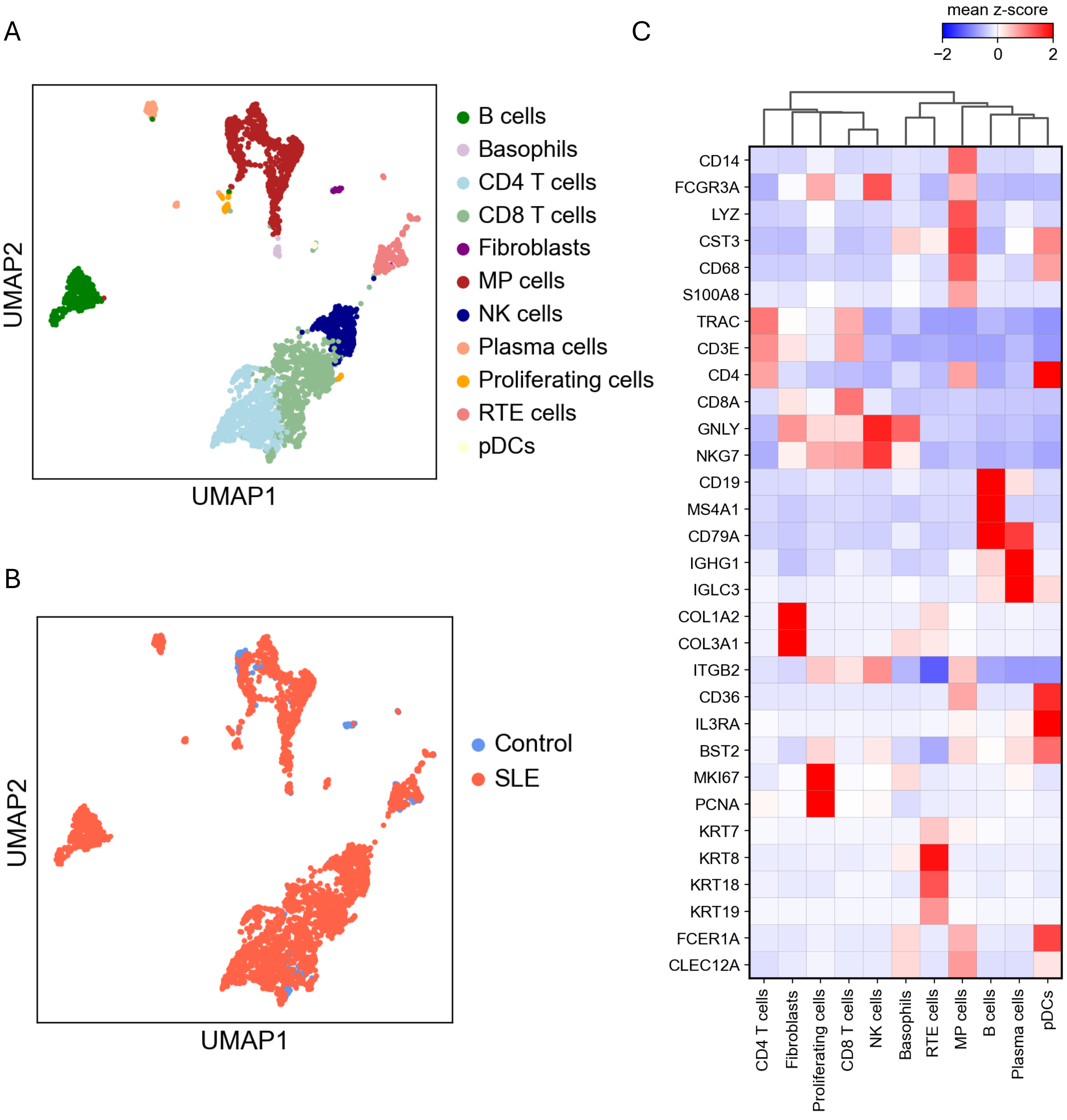


Supplementary figure 7. Single-cell profiling of lupus nephritis by scRNA-seq (data from the NIH Accelerated Medicine Program (AMP) for SLE (ImmPort, access code SDY997)). (A) UMAP visualization of cells assigned to 11 cell types based on their transcriptomic profiles (RTE = Renal Tubular Epithelial, MP = Mononuclear Phagocytic, pDCs = Plasmacytoid Dendritic cells). (B) UMAP colored by condition (blue = healthy control kidney, red = lupus nephritis kidney). (C) Scaled heatmap of mean expression of select DEGs and markers used for cell type annotation.

### Supplementary Figure 8

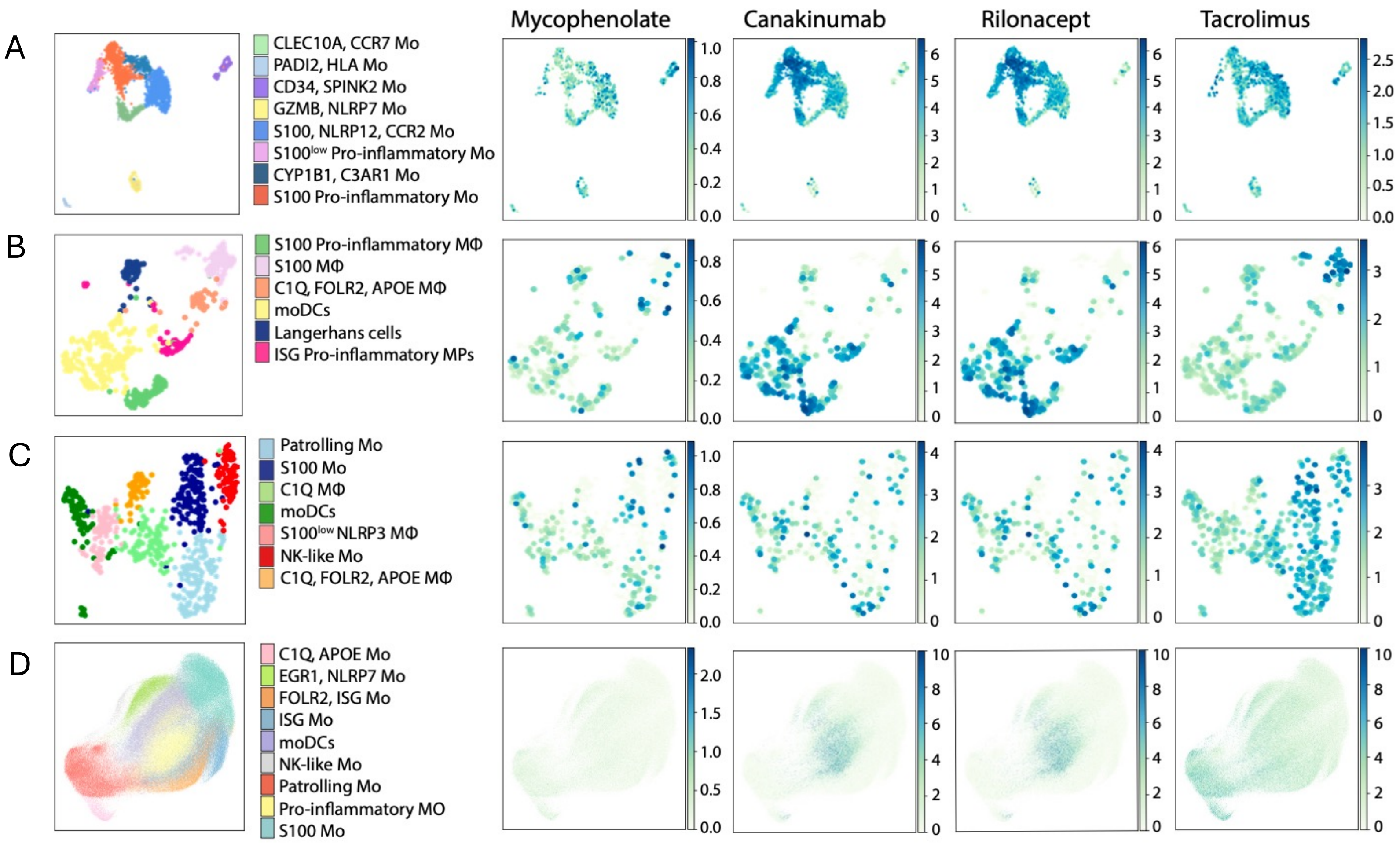


Supplementary figure 8. Single-cell drug prediction analysis. (A-D) Feature plots of selected predicted drug-target interactions based on target gene expression in (A) monocytes (Mo) stimulated by snRNP IC, (B) MPs in acute cutaneous lupus, (C) MPs in lupus nephritis, and (D) circulating Mo in SLE (Data from Figures 1, 4, and 5, respectively)
