## Supplementary Materials and Methods for "Lupus Immune Complexes Drive Distinct Pro-Inflammatory Monocyte and Macrophage Populations Independent of Type I Interferon"

### ***Processing, Analysis and Visualization of scRNA-seq data from snRNP immune complex (IC) stimulated monocytes (Mo)***

The raw scRNA-seq data were processed and aligned to the human reference genome, hg38, using Cell Ranger (v7.1.0, 10x Genomics) *(1)*. The filtered gene-cell barcode count matrices were further analyzed using the R (version 4.1.3) *(2)* based Seurat (version 4.2.1) package *(3)*. Pre-processing, normalization, scaling, dimensionality reduction and clustering were performed following the standard Seurat analysis pipelines as outlined below.

The cells were demultiplexed based on hashtag oligo (HTO) enrichment. Doublets and negatives were removed and singlets were extracted. Cells with fewer than 200 or more than 4,000 unique detected genes were filtered out as were cells with >5% mitochondrial counts. 3,040 Mo remained after filtering and were used for downstream analysis. Feature expression measurements for each cell were normalized by total expression and log-transformed. The top 2,000 highly variable genes were then identified. The expression levels were scaled so that the mean expression of each gene across cells is 0 and the variance is 1.

Linear dimensionality reduction was performed with Principal Component Analysis (PCA), and the first 30 principal components were used for downstream analysis. Non-linear dimensionality reduction was performed using Uniform Manifold Approximation and Projection (UMAP). The cells were embedded in a K-nearest neighbor (KNN) graph constructed using Euclidean distance in PCA space. The graph was then refined into a shared nearest neighbor (SNN) graph by weighting edges based on overlap of local neighborhoods, as implemented in the FindNeighbors() function. The Louvain *(4)* community detection algorithm was employed to determine the clustering of cells using the FindClusters() function with resolution = 0.4. Clusters were annotated using differentially expressed genes (DEGs), known canonical markers and literature review.

Differential gene expression analysis was performed using the FindMarkers() function, which identifies DEGs using the non-parametric Wilcoxon rank-sum test, with *p*-values adjusted for multiple testing using Bonferroni correction *(5)*. DEGs were visualized using EnhancedVolcano (version 1.12.0) *(6)*. Relevant biological processes and gene-gene correlations were evaluated using Gene Set Enrichment Analysis (GSEA) *(7)*. GSEA with Gene Ontology Biological Processes (GO BP) and KEGG (Kyoto Encyclopedia of Genes and Genomes) pathway gene sets were performed using clusterProfiler (version 4.2.2) *(8)*. Trajectory analysis was performed using the Python-based scFates (version 1.0.6) package *(9)* following standard pipelines. Transcription factor activity inference and pathway enrichment analyses were performed using the univariate linear model (ULM) method implemented in the Python-based decoupler (version 2.1.2) package *(10)*. Drug-target interactions were predicted using Drug2Cell (version 0.1.1) which determines interactions based on expression of target genes *(11)*.

Data visualization and downstream analyses were performed using plotting functions within Seurat along with ggplot2 (version 3.5.2) *(12)*. Plots were modified using the RColorBrewer (version 1.1-3) *(13)* and viridis (version 0.6.5) *(14)* packages. Stacked barplots were generated using dittoSeq (version 1.14.2) *(15)*. Additional feature plots and heatmaps were generated using the Python-based (version 3.10.12) *(16)* Scanpy (version 1.9.3) package *(17)*.

***Microarray data processing and differential expression analysis***

Raw expression data were normalized using the quantile method *(18)* and batch effect was corrected using the ComBat method *(19)*. The batch effect was assessed using PCA sample plot. Differential expression analysis was performed using the integrated hypothesis testing method *(20)*. Briefly, lupus immune complex (IC) treated and untreated groups were compared through empirical T-test and median difference test. The empirical null distributions were generated by 1,000 random permutations of the samples. Given the null distributions for T statistics and median differences, we computed two p-values against actual T-statistics and median difference. Stouffer's method was applied to compute combined p-value. False discovery rate (FDR) was estimated by Storey's method *(21)*. Finally, DEGs were selected with FDR < 0.05 and |log2-fold-change| ≥ 1. Three independent comparisons for each lupus IC were performed and identified each set of DEGs. We then compared the three sets of DEGs for up- and down-regulation, separately, resulting in overlapping and unique DEGs. With union of DEGs from the three comparisons, hierarchical clustering with cosine distance and ward linkage was performed to identify the differential expression patterns. The correlation analyses of log2-fold-change of the DEGs from each pair of the three lupus IC were performed using Spearman's method. Functional enrichment analysis of the DEGs was performed using DAVID *(22, 23)*, significantly enriched KEGG pathways were selected with p-value<0.01.

***SomaScan® Proteomic Analysis of Culture Supernatants of Immune Complex stimulated Mo***

Normalization of the protein levels from the Somascan® assay was performed by adaptive normalization by maximum likelihood per SomaLogic standard protocol *(24)*. The resulting ADAT file was read using the R based SomaDataIO package (version 6.1.0). Analyte and metadata information were appended to the ADAT data. The data were log-transformed after which differential protein expression analysis between the stimulated and unstimulated groups was performed using limma (version 3.58.1) *(25)*, with *p* values adjusted for multiple comparisons using the Benjamini-Hochberg method *(26)*. GSEA was performed using clusterProfiler (version 4.2.2) *(8)*. Spearman correlation analysis between upregulated (log2-fold-change > 0.5) genes from microarray analysis and upregulated proteins (log2-fold-change > 0.5) was performed.

A gene set comprised of genes corresponding to upregulated differentially expressed proteins (log2-fold-change > 2) in response to snRNP IC stimulation was created and labeled 'snRNP IC Upregulated Protein Signature'.

### *Analysis of SLE PBMC scRNA-seq data*

Single cell RNA sequencing data from peripheral blood mononuclear cells (PBMCs) in SLE (GSE174188) *(27)* were acquired from the NCBI Gene Expression Omnibus data repository and interrogated. This data is comprised of scRNA sequencing of PBMCs from 162 lupus patients ('SLE' group) and 99 matched healthy controls ('Healthy' group). Platelets were filtered out by removing cells with expression level >1 of platelet marker genes (*PF4, PPBP*). Normalization, scaling, dimensionality reduction and clustering were performed as described above. Clusters with high expression of hemoglobin genes (*HBA1, HBA2, HBB*) were removed. A subset of myeloid cells, identified by clusters with expression of *LYZ, CD14,* and *FCGR3A,* was created and comprised of 240,115 cells. Downstream analysis including scaling, dimensionality reduction, clustering (resolution = 0.4), differential gene expression, and visualization were performed as in scRNA-seq analysis of snRNP IC-stimulated Mo.

Module scores for expression of the 'Gene Ontology Biological Process Response to Type I interferon', 'snRNP IC Gene Signature', 'snRNP IC Upregulated Protein Signature' and 'ETS2 associated genes' gene sets were calculated for each cluster using Seurat. The 'ETS2 associated genes' gene set was derived from genes associated with ETS2 overexpression as identified by Stankey et al *(28)*. Module scores for expression of upregulated genes (log2-fold-change > 2) identified by microarray analysis of Mo stimulated with snRNP IC, dsDNA IC, and Ro IC were also calculated for each cluster. Kruskal-Wallis testing was performed for comparison of module scores across cell clusters using ggpubr (version 0.6.0) *(29)*. Transcription factor activity inference and pathway enrichment analyses were performed using the univariate linear model (ULM) method implemented in decoupler (version 2.1.2) *(10)*. Drug-target interactions were predicted using Drug2cell (version 0.1.1) *(11)*.

### *Analysis of Cutaneous Lupus scRNA-seq data*

Publicly available scRNA-seq data of skin biopsies from active cutaneous lupus (GSE186476) *(30)* were acquired from the NCBI Gene Expression Omnibus data repository and interrogated. scRNA-seq data from 7 skin biopsies from healthy control donors (GSM5652699, GSM5652700, GSM5652701, GSM5652702, GSM5652703, GSM5652704, GSM5652705) and skin biopsies from 7 patients with active cutaneous lupus lesions (GSM5652713, GSM5652714, GSM5652715, GSM5652716, GSM5652717, GSM5652718, GSM5652719) were pre-processed, normalized and integrated using anchor-based canonical correlation analysis *(31)* in Seurat. Scaling, dimensionality reduction, and clustering (resolution = 0.3) were performed as described above and the resultant clusters were annotated using canonical cell type markers and differentially expressed genes. A subset of myeloid cells, identified by expression of *LYZ, CD14,* and *FCGR3A,* was created and comprised of 591 cells. Downstream analysis including scaling, dimensionality reduction, clustering (resolution = 0.4), differential gene expression, GSEA, transcription factor activity inference, pathway activity inference analyses and visualization were performed as outlined above.

Module scores for expression of the ‘Gene Ontology Biological Process Response to Type I interferon’, ‘snRNP IC Gene Signature’, ‘snRNP IC Upregulated Protein Signature’ and ‘ETS2 associated genes’ gene sets were calculated for each cluster using Seurat. Module scores for expression of upregulated genes identified by microarray analysis of Mo stimulated with snRNP IC, dsDNA IC, and Ro IC were also calculated for each cluster. Kruskal-Wallis testing was used for comparison of module scores across cell clusters.

Cell-cell communication inferential analysis was performed using CellChat (version 2.1.2) *(32)*. Comparative ligand-receptor interaction analysis was performed between healthy control skin and cutaneous lupus skin. Transcription factor activity inference and pathway enrichment analyses were performed using the univariate linear model (ULM) method implemented in decoupler (version 2.1.2) *(10)*. Drug-target interactions were predicted using Drug2cell (version 0.1.1) *(11)*.

***Analysis of Lupus Nephritis scRNA-seq data***

Single cell RNA sequencing data of kidney biopsies from patients with lupus nephritis were acquired through the NIH AMP *(33)* SLE program for analysis. This data is comprised of scRNA sequencing using the CEL-Seq2 protocol performed on kidney biopsies from 24 patients with lupus nephritis (class III and IV) and 10 control samples (living donor kidney biopsies) totaling 2,736 immune cells and 145 renal epithelial cells. The data were pre-processed, normalized and integrated using anchor-based canonical correlation analysis *(31)* in Seurat. Scaling, dimensionality reduction, and clustering (resolution = 0.8) were performed as described above and the resultant clusters were annotated using canonical cell type markers and differentially expressed genes. A subset of myeloid cells, identified by expression of *LYZ, CD14,* and *FCGR3A,* was created and comprised of 687 cells. Downstream analysis including scaling, dimensionality reduction, clustering (resolution = 0.8), differential gene expression, GSEA, transcription factor activity inference, pathway activity inference and visualization were performed as outlined above. Inferential analysis of cell-cell communication via ligand receptor interaction enrichment was performed using CellChat (version 2.1.2) *(32)*. Transcription factor activity inference and pathway enrichment analyses were performed using the univariate linear model (ULM) method implemented in decoupler (version 2.1.2) *(10)*. Drug-target interactions were predicted using Drug2cell (version 0.1.1) *(11)*.

Module scores for expression of the ‘Gene Ontology Biological Process Response to Type I interferon’, ‘snRNP IC Gene Signature’, ‘snRNP IC Upregulated Protein Signature’ and ‘ETS2 associated genes’ gene sets were calculated for each cluster using Seurat. Module scores for expression of upregulated genes identified by microarray analysis of Mo stimulated with snRNP IC, dsDNA IC, and Ro IC were also calculated for each cluster. Kruskal-Wallis testing was used for comparison of module scores across cell clusters.

***Western Blot***

For protein analysis, human Mo freshly isolated from PBMCs were stimulated with snRNP ICs for 2 hours. Protein extracts were separated by SDS-PAGE and transferred onto PVDF membranes. Membranes were probed with primary antibodies against total ETS2 and phospho-ETS2 (Thr72) (Invitrogen), and β-actin (Santa Cruz Biotechnology) as a loading control. After washing, membranes were incubated with HRP-conjugated secondary antibodies (Santa Cruz Biotechnology). Protein bands were visualized using Pierce ECL Western Blotting Substrate (Thermo Scientific) and detected by chemiluminescence.

***MEK inhibition***

Human Mo freshly isolated from PBMCs were pretreated with the MEK inhibitor PD0325901 (mirdametinib; MedChemExpress, LLC) for 30 minutes, followed by stimulation with snRNP ICs. For transcriptomic analysis, cells were stimulated for 4 hours, after which total RNA was extracted and used to generate cDNA and barcoded sequencing libraries with the TruSeq Stranded mRNA Kit (Illumina) according to the manufacturer’s instructions. Libraries were sequenced on an Illumina HiSeq 4000 platform using a 2 × 100 bp paired-end configuration. For cytokine analysis, Mo were stimulated for 12 hours, and levels of IL-1β and IL-6 in culture supernatants were measured by ELISA. Normality was assessed using the Shapiro-Wilk test. A paired t-test was used to compare cytokine levels in the presence and absence of MEK1/2 inhibitor.

### *Analysis of bulk RNA-seq data*

Pseudoalignment-based transcript quantification of the raw sequencing data was performed using Salmon (v1.10.3) *(34)* with the *Homo sapiens* GRCm38 reference transcriptome (https://www.gencodegenes.org/human/). The resulting transcript-level abundance estimates were imported into R (v4.5.1) and aggregated to gene-level counts using tximport (v 1.38.2) *(35)*. Differential expression analysis was conducted with DESeq2 (v1.50.2) *(36)* using the default Wald test. P-values were adjusted for multiple testing using the Benjamini-Hochberg method *(26)*. Differentially expressed genes with adjusted p-value < 0.05 were considered significant. Heatmaps of gene expression were generated from variance-stabilized transformed (VST) counts. Transcription factor activity inference was performed using the univariate linear modeling (ULM) method implemented with the Python (v3.11.14) version of decoupler (v2.1.2) *(10)* and the CollecTRI database. Gene set enrichment analysis (GSEA) *(7)* was performed using clusterProfiler (v4.18.4) *(8)* with Kyoto Encyclopedia of Genes and Genomes (KEGG) pathway gene sets, using a ranked list of genes based on DESeq2 Wald statistics. Gene sets with a false discovery rate (FDR) < 0.05 were considered significantly enriched.
